# A microbiota-dependent fitness locus promotes intraspecies competition and cell envelope resilience in gut bacterium *Phocaeicola vulgatus*

**DOI:** 10.64898/2026.11.24.768882

**Authors:** Yelizaveta Rassadkina, Jayanth Jawahar, Amina Bradley, Rachel L. Neve, Per N. Malkus, Erica J. Washington, Elizabeth A. Shank, Jessica R. McCann, John F. Rawls

## Abstract

A central question in gut microbial ecology is how dominant bacterial species establish and occupy their niches amid inter- and intraspecies competition. *Phocaeicola vulgatus* (*Pvu*) is a Gram-negative anaerobe and prevalent member of the human gut microbiome that can integrate into established gut communities without antibiotic pretreatment, yet mechanisms underlying this competitive success remain poorly understood. Using *in vivo* transposon insertion sequencing, we identified a three-gene locus that promotes *Pvu* intraspecies fitness in the presence of a complex microbiota but is dispensable in germ-free hosts. The locus encodes a predicted HU/IHF superfamily histone-like DNA-binding protein (DifH), a hypothetical lipid-binding protein (DifS), and a FadL-like outer-membrane transporter (DifT). *In vitro* genetic and transcriptomic analyses implicated these genes in cell envelope resilience. Chromatin immunoprecipitation sequencing revealed that DifH binds a specific repetitive DNA motif at its own locus and a limited set of additional loci involved in metabolism, envelope biogenesis, and cellular defense. Together, these findings identify a previously unrecognized fitness locus that promotes cell envelope resilience, enables intraspecies competition, and supports *Pvu* persistence within complex gut microbial communities. More broadly, these results uncover mechanisms by which this dominant gut commensal secures its niche and suggests avenues to therapeutically modulate gut microbiota composition.

## INTRODUCTION

The mammalian gastrointestinal tract represents one of the most complex microbial ecosystems, harboring hundreds of bacterial species that are organized into spatially and metabolically distinct communities^1,2^. Bacterial colonization and persistence within this environment are shaped by numerous physiological gradients, including oxygen availability, antimicrobial exposure, pH, nutrient availability, mucus composition, and intestinal transit rate^3,4^. These heterogeneous conditions create specialized ecological niches that select for bacteria capable of adapting to those local environments and engaging in complex networks of inter- and intraspecies interactions^3,5,6^. This diversity of microenvironments also enables different strains of the same bacterial species to coexist within a single host^7,8^. Despite extensive characterization of gut microbial community composition, the principles and mechanisms that enable individual bacterial species to persist, compete, and adapt within these complex communities remain very poorly understood.

Members of the *Bacteroidetes* phylum represent some of the most abundant and prevalent organisms in the mammalian gut. However, functional characterization has largely focused on a limited number of well-studied species including *Bacteroides thetaiotaomicron* and *Bacteroides fragilis*^9,10^. The focus on these strains has left other related species, such as *Phocaeicola vulgatus* (hereafter referred to as *Pvu*; formerly *Bacteroides vulgatus*^11^) largely understudied. *Pvu* is a highly prevalent member of the human intestinal microbiota that is acquired early in life^12–15^, and is one of the most globally dispersed human gut bacterial species^16^. *Pvu* also readily colonizes established complex microbial communities^8,17–19^, and has been identified as a gatekeeper that can inhibit colonization of other bacterial species, including after fecal microbiota transplant^20^. *Pvu’*s recent reclassification from the *Bacteroides* to the *Phocaeicola* genus underscores its evolutionary divergence from more extensively studied *Bacteroides* species^11^. Thus understanding the genetic determinants that underlie the successful colonization and persistence of this prevalent human commensal within the gut ecosystem is an important research goal.

Transposon insertion sequencing (Tn-seq; also called insertion sequencing, IN-seq) has been used to identify genes involved in diverse biological processes across the *Bacteroidetes* (including *Bacteroides, Phocaeicola,* and *Parabacteroides* species) revealing genes involved in nutrient acquisition and metabolism^10,21–24^, bacterial responses to environmental stresses and signals^23–26^, and those involved in morphology and survival^27,28^. Here, we used a genome-wide Tn-seq approach to identify genetic determinants required for *Pvu* fitness during colonization of germ-free (GF) mice and those reared in the presence of a complex microbiota (specific pathogen-free; SPF). We report a previously uncharacterized locus that promotes intraspecies *Pvu* fitness specifically within the competitive SPF environment. Through *in vitro* assays, transcriptomic, and DNA-binding analyses, we reveal that the genes in this locus contribute to bacterial cell envelope resilience, representing a previously unrecognized fitness system that enables *Pvu* to compete with its own species to persist in the ecological context of a complex gut microbiota.

## MATERIALS AND METHODS

### *In vitro* bacterial culture

All *Pvu* strains were grown in a flexible anaerobic chamber (Coy Laboratory Products) with an atmospheric composition of 5% H_2_, 5% CO_2_, and 90% N_2_. Overnight cultures prepared for gavage were grown in liquid BHIS medium (BHI; BD Bacto, #237500, supplemented with hemin (final concentration, 2.5 µg/mL) and vitamin K (final concentration, 0.5 µg/mL) using a 1:100 dilution of the commercial mixture (Hardy Diagnostics, #Z237). Experiments with minimal medium were performed using *Bacteroides* Minimal Medium (BMM) supplemented with 0.5% glucose as the sole carbon source (Sigma, # G7528)^29^.

For growth curve assays, cells were inoculated 1:100 from an overnight culture into fresh, pre-reduced liquid medium, and grown in triplicate in 96 well non-treated, flat-bottomed tissue culture plates (VWR, 734-2781). For the cell membrane susceptibility assays, polymyxin B (Sigma, #P4932), carbenicillin (Fisher, #10177012), meropenem (Sigma, #ATE517248713), fosfomycin (Sigma, #P5396), sodium deoxycholate (Sigma, #30970), bile salts (Sigma, #48305), TCA (Sigma, #T4009), and cholic acid (Sigma, #C1129) were add in concentrations as stated in figure legends. Approximately 50µL of mineral oil (Sigma, #330779) was added to the top of culture plates to prevent evaporation of the growth medium for the duration of the assay. OD600 was measured once per using the Spectrostar Nano (BMG Labtech). Growth curves were then analyzed and graphed using GraphPad Prism version 11.0.2.

### Transposon mutagenesis of *Pvu*

Transposon mutagenesis was performed via conjugative transfer of the pSAM plasmid (Addgene, Cat# 102939) as previously described for *Bacteroides thetaiotaomicron*^30^ with some modifications. Briefly, the aerobic incubation of *E. coli* with *Pvu* was extended to 8 hours. Approximately 45,000 colonies were pooled in TYG media (Sigma, #MB000000T) supplemented with 20% glycerol (Fischer, Cat# BP229-1) and stored at -80°C in 1mL aliquots. Aliquots were used directly for arraying and combinatorial pooling to generate the arrayed *Pvu* Tn library. Outgrowths of the aliquots were done to obtain enough bacteria for GF and SPF mouse gavages. 1mL aliquots were thawed at room temperature under anaerobic conditions and added to 50mL of pre-reduced TYGs medium supplemented with 5µg/mL erythromycin (Sigma, #E5389), 50µg/mL gentamicin (Sigma, #G1264), and 0.5g/L cysteine (Sigma, #168149). The cells were incubated at 37°C for 24h, during which O.D.600 remained unchanged. After 24 hours, the cells were incubated at room temperature for an additional 24 hours, after which the O.D.600 was 1.19. This growth at 25°C was performed to not bias the Tn library towards strains with a faster replication rate. 22mL of 50% glycerol was added to ∼50mL of the outgrowth Tn library for a final glycerol concentration of 15% and 2mL aliquots were stored at -80°C. Library aliquots were then plated aerobically and anaerobically on TSA blood agar (VWR, #10324-332) plates to test for sterility as well as to count CFUs, which were found to be approximately 9.2x10^8^ CFU/mL. SPF samples were pooled per cage to get sufficient material for Tn-seq. Tn-seq analysis of outgrowth replicates compared to the original transduced library showed no significant differences in Tn mutant representation (data not shown).

### Tn-seq analysis of transposon mutant populations

For the combinatorially pooled library, raw FASTQ reads were mapped to the genome assembly of *Pvu* CL09T03C04, using the genome annotation generated in this study. Count tables were generated for each pool, and well and plate assignments were determined for each mutant using previously described protocols^31^. For the input and output Tn-seq libraries obtained from *in vivo* experiments, the raw fastq reads were analyzed using the TRANSIT resampling^32^ tool as implemented on the BV-BRC bioinformatics database^33^ (Supplementary Table S3). For PCoA, insertion-level read counts were filtered to TA sites with at least 3 reads in at least 2 of the 19 samples (10%), converted to presence/absence, and compared using Jaccard (binary) distance. PCoA, PERMANOVA (adonis2, 9,999 permutations), and tests of multivariate dispersion (betadisper) were performed using a fixed random seed. Because multiple samples were collected from each mouse, the effect of week was tested with permutations restricted within mouse, and the effect of microbiota status was tested on one terminal sample per mouse. Statistics were generated using the vegan package^34^ (v2.7-2) in R^35^ (v4.6.0), while the PCoA and barplots were generated using the ggplot2^36^ (v4.0.3) and the heatmaps with pheatmap^37^ (v1.0.13).

### Protein function prediction

Alphafold^38^ was used to predict the structures of the *dif* locus. Predicted structures were evaluated based on model confidence scores and visual inspection of the resulting structural models. For DifS (Pvu2207), AlphaFold predicted a protein structure containing substantial intrinsically disordered regions, resulting in lower confidence across portions of the model. To further assess protein disorder, the amino acid sequence of DifS was analyzed using PONDR^39^ and IUPred3^40^. Structural similarity searches were performed using FoldSeek^41^ to identify proteins with related structural features. The predicted folded core of DifS was visualized using UCSF Chimerax^42^. Then, to predict protein function, InterPro^43^, CDART^44^, and eggNOG^45^ were used to obtain predicted functions, signal sequences, and conserved domains. For DifH and its close relatives, InterPro^43^ and HHPred^46,47^ were used to identify the HU-HIG and the winged helix-like superfamily domains.

### Generation of PvuCL09difS-FLAG

We generated the FLAG-tagged strain by introducing a sequence encoding *difS* fused to a FLAG tag, followed by a stop codon, into a tRNA integration site in the *ΔdifS* knockout strain following an established protocol^48^. *PvuCL09fifS-FLAG* strains were then validated as described for the clean deletion strains (Supplemental Methods). Expression of *PvuCL09difS -FLAG* was confirmed by western blot using the following protocol: *Pvu* strains were grown to an O.D.600 of 0.6, pelleted, resuspended in 250uL of RIPA buffer (Sigma, #R0278) with protease inhibitor cocktail (Roche, #11836145001), and rocked for 30 minutes at 4°C. The samples were then spun down at 2 min at 10,000xg and the supernatant was loaded into wells of a 10% mini-protean® tgx™ precast protein gel (BioRad, #4561035). Proteins were then transferred onto a nitrocellulose membrane (BioRad, # 1620112) using the Trans-blot Turbo Transfer System (Biorad). The membrane was then blocked with 5% bovine serum albumin (BSA; Fisher, #BP1600) in Tris-Buffered Saline (TBS; 200 mM Tris Base, Fisher Cat# BP152-500, and 1500 mM NaCl, Sigma Cat# S9888, and pH adjusted to 7.6), with 1mL Tween 20 (Sigma, Cat#P1379) (TBST) buffer for one hour at room temperature. The resulting membrane was treated with anti-FLAG tag antibody (Cell Signaling, #8146) diluted at 1:1000 in 5% BSA in TBST and incubated overnight at 4°C, washed 3x with TBST for 5 minutes, and anti-FLAG was detected with secondary horse anti-mouse HRP linked antibody (Cell Signaling, #7076) 1:2000 in 5% BSA in TBST, incubated for one hour at room temperature. The membrane was washed again 3x with TBST for 5 minutes followed by addition of Clarity™ Western ECL Substrate (BioRad, #1705060) to then visualize the proteins using a Chemidoc system. Kaleidoscope protein ladder (BioRad, #1610375) was used to identify *PvuCL09difS-FLAG*.

### Animal husbandry

All animal procedures were performed at Duke University (AAALAC Accreditation Number 363) according to protocols approved by the Duke University Institutional Animal Care and Use Committee (IACUC; protocol registry numbers A111-23-05 and 103). This protocol was prepared and registered at Duke University before the study occurred.

#### Specific pathogen-free (SPF) mice

All animal experiments were done at Duke University according to IACUC-approved protocols. C57BL/6J male and female SPF mice at eight weeks of age were obtained from Jackson laboratories (Strain #000664). Upon arrival at Duke, mice were allowed to acclimate to the facility for one week and housed in cages containing Enrich-o’Cobs bedding (The Andersons Lab) for environmental enrichment. Each cage housed 3-5 mice in an ABSL2 animal facility with ad libitum access to food (PicoLab Rodent Diet 20 #5R53; LabDiet) and water. Then, fecal pellets were collected from each cage to confirm that the mice did not harbor endogenous *Pvu* via PCR with *Pvu* specific primers^30^ (Supplementary Table 7). Subsequently, mice were gavaged at eight weeks of age, by trained personnel to minimize handling stress, with 200µL of *Pvu* bacterial cells (∼10^8^-10^9^ cells/mL) washed in pre-reduced phosphate-buffered saline (PBS). Since these procedures were not expected to cause more than momentary discomfort, no anesthesia or analgesia was administered. Mice were weighed at gavage and at each subsequent weekly fecal collection time point. Mice were monitored daily. At 4 weeks post-gavage, mice were euthanized by CO₂ asphyxiation followed by bilateral thoracotomy, and fecal samples, tissues, and cecal contents were collected and immediately snap-frozen in liquid nitrogen. A total of 66 SPF mice were used across all experiments in this study.

#### Germ-free (GF) mice

The mice were housed in cages of 2-4 mice in a flexible film gnotobiotic isolator (Class Biologically Clean) with ad libitum access to both food (2020SX Teklad diet; Envigo) and water. Cages were provided with ALPHA-dri® bedding (Shepherd Specialty Papers, Inc.) for environmental enrichment. Both male and female germ-free C57BL/6N mice were gavaged at eight weeks of age as described for SPF animals and were monitored daily. At four weeks post-gavage, mice were taken out of the gnotobiotic isolator and transferred immediately to a biological safety hood for euthanasia by CO₂ asphyxiation followed by bilateral thoracotomy. Fecal samples, tissues, and cecal contents were removed and immediately snap frozen in liquid nitrogen. A total of 21 GF mice were used across all experiments in this study.

#### Humane endpoints and animal monitoring

Mice were monitored daily and weighed weekly throughout the experiment. Animals were assessed for clinical signs of distress, including hunched posture, ruffled fur, reduced activity, diarrhea, and dehydration. No mice reached the humane endpoint of 20% weight loss or clinical signs of distress as described above during the study.

### qPCR analysis of *in vivo* competition

DNA was extracted from fecal samples (Zymo Quick-DNA fecal/soil microbe miniprep kit Zymo, #D6010) and subjected to qPCR with primers specific for all *Pvu*, *Pvu dif* deletion specific, and universal 16S rRNA genes (Supplementary Table 7). For SPF mice, all three primer sets were used to obtain a relative percent abundance of WT *Pvu* and each *dif* deletion strain as compared to total 16SrRNA gene content, while in GF only Pvu- and deletion-specific primers were used. For the SPF mice, the percent relative abundance was calculated by first generating a coefficient using the efficiency percent of each primer set as determined by its standard curve. This coefficient was then used in the formula: (coefficient for 16S rRNA primers^(Avg Cq value of 16S primers for sample)^/coefficient for Pvu primers^(average Cq value of Pvu primers for sample)^)*100, resulting in relative abundance of *Pvu* as a % of total bacteria present^49^. To obtain the individual percent of each strain, the above equation was used again to define the percentage of each deletion with deletion specific primers, then that value was subtracted from the total *Pvu* percent to obtain the WT *Pvu* percent. For GF mice, the same equation was used with total *Pvu* determined by *Pvu* specific primers, and percentage of deletion determined with deletion specific primers. All samples were run in duplicate on the TB Green® Premix Ex Taq™ II FAST qPCR (Takara, Cat# RR830B) (Supplementary Table S7). These data were then used to plot the percent relative abundance of each strain as well as calculate the deletion/WT ratios to create a competitive index using GraphPad Prism version 11.0.2. For mice given only a single *Pvu* strain, comparisons between WT and deletion strains at each time point were performed using the Mann-Whitney test. For mice co-gavaged with WT and deletion strains, a Student’s *t*-test was used to compare the deletion/WT abundance ratio at each time point with the input gavage ratio.

### Competition across intestinal sections

Mice (n = 6) were euthanized and the intestine was collected from the duodenum to the rectum. The small and large intestines were cut from the cecum and cross sectioned every 2cm. Sections were flayed open and placed into wells of 24 well plate (Eppendorf, #30722019) with 500mL of sterile ice-cold 1x PBS (Corning, Cat#46-013-CM). Samples were shaken for 10 minutes at 4°C, and resulting well contents were moved to 1.5mL Eppendorf tubes and frozen at -80°C. DNA from each sample was extracted (Zymo Quick-DNA fecal/soil microbe miniprep kit #D6010) and processed as described for fecal sample DNA.

### Generation of *PvuCL09difH-HA*

We inserted sequence coding for an in-frame HA tag just upstream of the native DifH stop codon. This insertion was generated in the same manner as the clean deletions (Supplemental Methods) following an established protocol^48^. *PvuCL09difH-HA* strains were validated as described above for clean deletion strains. Expression of *PvuCL09difH-HA* was confirmed by western blot using a similar protocol as described for the *PvuCL09difS-FLAG,* with the following modifications: Resuspended pellets were sonicated using a Branson Ultrasonics Tip Sonifier for a total of 2 minutes with 30 millisecond pulses and 15 second rest periods on ice. The sonicated cells were then spun down for 5 min at 10,000 x g at 4°C and the supernatant was collected. Soluble protein was quantified using the Pierce BCA assay kit (Thermo #23227) according to manufacturer’s instructions. Then 80ug of each sample were loaded into each well of a 10% mini-protean® tgx™ precast protein gel (BioRad, #4561035). Proteins were then transferred onto a nitrocellulose membrane (BioRad, #1620112) using the Trans-blot Turbo Transfer System (Biorad). The membrane was blocked with 5% bovine serum albumin (BSA; Fisher, #BP1600) in Tris-Buffered Saline with Tween 20 (TBST) buffer for 1 h at room temperature and then treated with anti-HA tag antibody (abcam, #ab9110) diluted at 1:1000 in 5% BSA in TBST and incubated overnight at 4°C, washed 3x with TBST for 5 minutes. An anti-HA was detected with secondary mouse anti-rabbit HRP linked antibody (Cell Signaling, #5127S) 1:2000 in 5% BSA in TBST, incubated for one hour at room temperature. The membrane was washed again 3x with TBST for 5 minutes followed by addition of Clarity™ Western ECL Substrate (BioRad, #1705060) to then visualize the proteins using a Chemidoc system.

### *In vitro* chromatin immunoprecipitation sequencing (ChIP-seq)

*PvuCL09difH-HA* and WT *Pvu* grown in BHIS and BMM following an adapted version of a previously established protocol^50^. 300mL of each strain was grown in BHIS (n = 4) and 500mL in BMM (n = 4), until O.D.600 of 0.6 and then formaldehyde treated (Fisher, #F79-1) with a final concentration of 1% of total volume at room temperature for 20 min. Then, 2M glycine (Fisher, #BP381-1) was added to each 50mL conical to a final concentration of 0.5M. Cells were then pelleted by spinning for 15 min at 3,220 x g at 4°C, washed in 2x TBS and resuspended in 1.5mL of lysis buffer as described^50^ in a 15mL conical and supplemented with 4mg/mL of lysozyme (Sigma, #L4919). Cells were incubated at 37°C for 30 minutes followed by a 5 min on ice and then sonicated using a Branson Ultrasonics Tip Sonifier in an ice water bath for 8 cycles with a 25 second sonication with 2 minute rests in between on 40% power. Sonicated samples were then transferred to a 1.5mL Eppendorf tube and spun at max speed for 15 minutes at 4°C. Supernatants were transferred into a fresh tube and 50uL was pooled from each sample for the input control. 10uL of each sonicated sample was visualized on a 2% agarose gel to confirm that sonication resulted in DNA shearing to a size range of 500bp. All samples were then stored at -80°C until immunoprecipitation.

For immunoprecipitation, 60μL of protein A+G magnetic beads (Sigma, #16-663) per sample were washed 4 times with 1mL of 1x PBS + 0.02% Tween-20 (Fisher, #BP337) at 4°C for 5 minutes with agitation. After the fourth wash, beads were resuspended in 250mL of 1x PBS + 0.02% Tween-20 and 8ug of the anti-HA tag antibody (Abcam, #ab9110) was added to each tube. The pre-incubated beads with the antibody and the lysate were transferred to a 2mL Eppendorf tube and rotated at 4°C overnight. Lysate was then discarded, and beads were washed in 1mL of 1x PBS + 0.02% Tween-20 with rotation for 5 minutes at 4°C followed by aspiration of the PBS. Input samples were thawed, and 100 µL of ChIP-seq elution buffer was added to all samples. Samples were then incubated at 65°C overnight to reverse cross-links.

Chip-seq DNA was then extracted by Phenol:Chrloroform:Isoamyl Alcohol (PCIA). Eluted samples were boiled at 95°C for 10 minutes. 300uL of H_2_O and 400uL of room temperature PCIA (Thermofisher, #15593049) was added to each tube. Samples were vortexed and transferred to phase lock tubes, spun at maximum speed at room temperature for 5 minutes and then the upper layer was transferred to new 1.5mL Eppendorf tube. 2uL of 20mg/mL glycogen (Thermofisher, #R0561), 40uL 3M sodium acetate (Corning, #46-033-CI), and 1mL 100% ethanol (Koptec, Cat# V1016) were added to each tube. Tubes were vortexed and incubated at -20°C for a minimum of four hours. Tubes were spun for 15 min at maximum speed at 4°C and supernatant was removed. 1mL of 70% ethanol was added per sample and samples were spun for 5 minutes at maximum speed at 4°C. Samples were dried and resuspended in 20uL of DEPC-treated water (Thermo, Cat# AM9915G). DsDNA HS Qubit kit (Invitrogen, #Q32851) was used to quantify DNA. All ChIP-seq samples were library prepped using the Watchmaker DNA Library Prep kit (WatchMaker, #7K0102) and sequenced on a NextSeq1000 XLEAP P1 with 300 cycles (150 x 150) to generate 100M paired end reads at the Duke Sequencing and Genomics Technology Core.

### ChIP-seq bioinformatic analysis

All fastq files of the sequenced samples were uploaded to and analyzed using the Galaxy web platform^51^. Reads were paired and then mapped against the *Pvu* CL09T03C04 genome using Bowtie2^52,53^ followed by bamCoverage^54^ to generate bigwig files of each sample. All bigwig files were then visualized individually and then as a log2ratio (*PvuCL09difH-HA*/WT or *PvuCL09difH-HA*/input) generated by bigwig compare^54^ using JBrowse2^55^ in alignment with *Pvu* CL09T03C04. Subsequently, MACS2^56,57^ and BEDtools ClosestBed^58^ were used to call peaks and identify their location across the genome by comparing all of the *PvuCL09difH-HA* samples to either the input or WT *Pvu* samples based on the genome size of *Pvu* CL09T03C04 and a q-value=0.0001. Once peaks were identified, their genomic coordinates were used alongside pooled replicate BAM files for the WT *Pvu* and *PvuCL09difH-HA* samples. Replicates were merged using Samtools^59,60^ merge and bamCoverage^54^. The resulting coverage data and peak coordinates were then used with computeMatrix^54^ to generate a matrix, which was visualized as a heatmap using plotHeatmap^54^. DNA-binding motifs of the BHIS and BMM samples were detected and visualized along the peaks using MEME^61,62^. All of the peak summits were then plotted using GraphPad Prism version 11.0.2. Detection of the binding motif along the peaks, the entire *Pvu* genome, DifH homologs, and its close congeners was done by FIMO^62^ on genomic regions obtained and visualized using BV-BRC^33^. The heatmap that combines the ChIP-seq peaks with the RNA-seq log2 fold change data was generated by pheatmap^37^ (v1.0.13).

### Supplemental methods

The Supplemental Methods document associated with this publication includes details on *Pvu* genome sequencing, sequencing arrayed *Pvu* Tn mutant library clones, RT-PCR of the *dif* locus, generation and validation of clean deletion strains, conservation analysis of *Pvu* and its close congeners, *in vitro* RNA-seq and analysis, LC-MS bacterial culture conditions, and metabolomic and lipidomic data processing.

## RESULTS

### Generation of a genome-wide *Pvu* transposon library to identify *in vivo* fitness determinants

To identify genetic determinants required for *Pvu* fitness, we generated a genome-wide transposon (Tn) mutant library in the *Pvu* strain CL09T03C04^63^, isolated from a healthy human, using established protocols^30^. The resulting pooled library contained approximately 40,000 clones representing close to 40,000 unique Tn insertion sites. This provided broad genomic coverage for unbiased identification of fitness-associated loci (Supplementary Tables S1 and S2). In parallel, we generated an arrayed collection of 1,100 individual mutant strains from the original library through combinatorial pooling, the majority of which contained a single Tn insertion, which enabled downstream characterization of individual mutants (Supplementary Figure S1A). Independent sequencing of separate aliquots from the constructed library^31,32^ demonstrated high reproducibility (Supplementary Figure S1B). Tn insertions were well distributed across the *Pvu* genome (Supplementary Figure S1C), confirming the quality and genome-wide coverage of the Tn mutant library.

To identify genes required for *Pvu* fitness during colonization of distinct microbial environments, the *Pvu* Tn library was gavaged into adult GF and SPF mice. Fecal samples were collected weekly over a four-week colonization period, followed by sacrifice and sample collection (Figure 1A). Genomic DNA was extracted from fecal and cecal samples for each time point and used for downstream analysis. All analyzed samples exhibited saturation of unique insertion sites with increasing sequencing depth, indicating that sequencing depth was sufficient to capture the majority of represented mutants (Supplementary Figure S1D). To quantify the fitness of individual Tn mutants during intestinal colonization, output-to-input abundance ratios were calculated using TRANSIT^32^ (Supplementary Table S3). Genome-wide comparison of mutant fitness profiles revealed that most insertion mutants exhibited similar enrichment or depletion patterns across both GF and SPF mice throughout the experiment. Thus most genes contributed similarly to fitness regardless of microbial context (Supplementary Figure S1E; Supplementary Table S3). When subsetting for Tn mutant significance in at least one GF and one SPF sample, we see that the Tn mutant representation for most of those significant genes is either increased or decreased across time and mouse status (Figure 1B). However, a small and distinct subset of mutants displayed divergent fitness profiles between GF and SPF mice, suggesting that those genes might specifically contribute to *Pvu* fitness in response to the microbial community rather than the host gut habitat alone (Figure 1C). Principal coordinates analysis (PCoA) showed that mutant library composition varied along two main axes: PCoA1 (23.5% of variance) tracked time after colonization, reflecting progressive loss of insertion diversity relative to the input library, whereas PCoA2 (13.9%) separated GF from SPF mice (Figure 1D). Time explained a significant fraction of the variation within mice (PERMANOVA, within-mouse permutations, R^2^ = 0.17, p = 0.0004), and GF and SPF mutant populations differed at the terminal time point (R^2^ = 0.24, p = 0.048, n = 7 mice; dispersion did not differ between groups, p = 0.46). The ordination was robust to equalizing read depth (Procrustes r = 0.998). Overall, these data suggest that both temporal adaptation and the presence of a complex microbial community dynamically shaped the composition of the *Pvu* Tn mutant pool.

**Figure 1:**
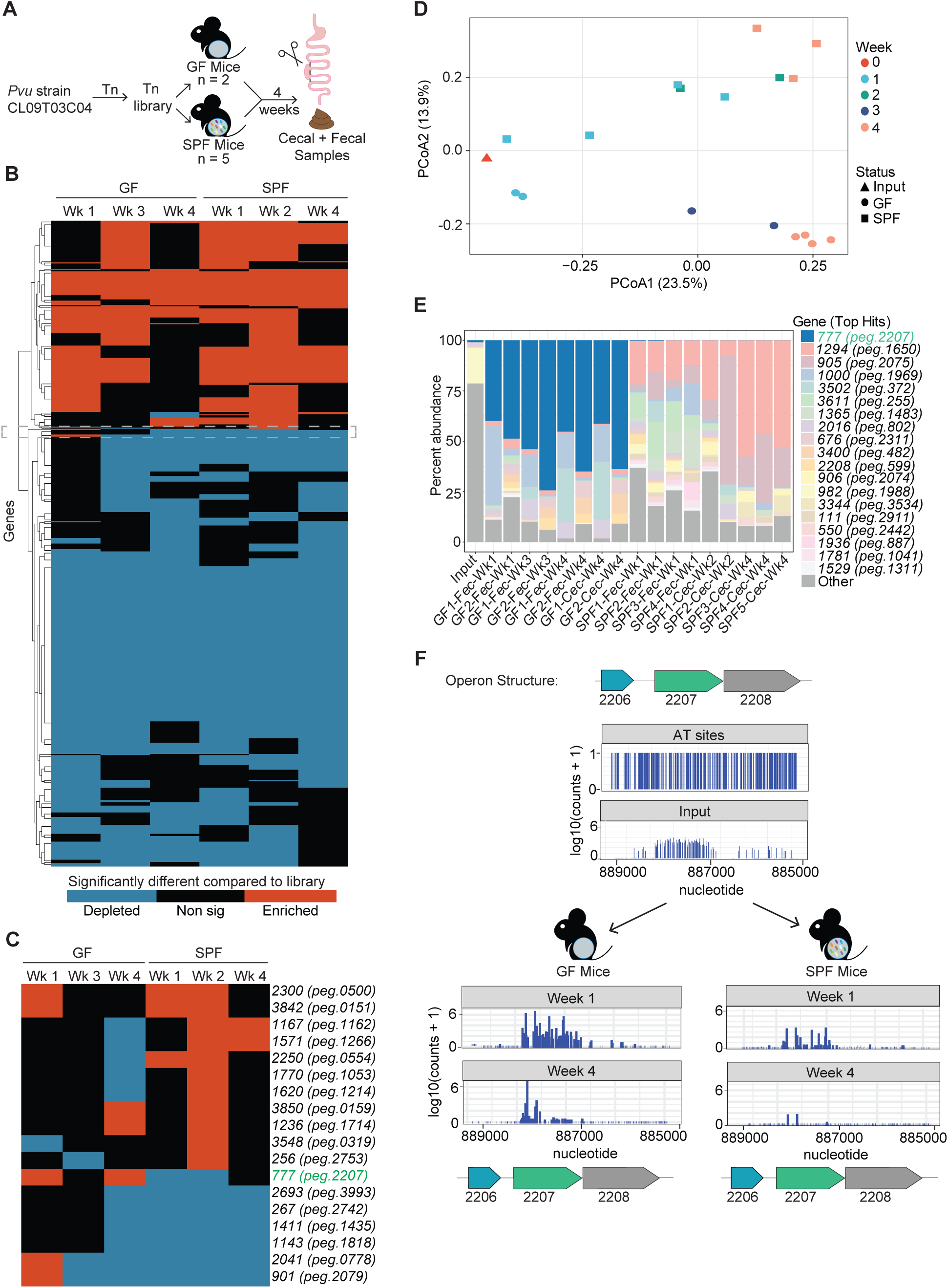
Tn-seq identifies *Pvu2207* as a context-dependent fitness determinant in the mouse intestine. **(A)** Schematic of the experimental design for the Tn-seq screen in GF (n = 2) and SPF mice (n = 5) **(B)** Heatmap of transposon mutants corresponding to genes that were significantly enriched or depleted by log2FC compared to WT (adjusted p < 0.05) in at least one GF and one SPF time point. A gray bracket highlights the genes shown in panel C. **(C)** Transposon mutants exhibiting discordant fitness phenotypes between the mouse colonization states, from panel B. **(D)** Principal coordinates analysis (PCoA) of all input and *in vivo* samples (n = 19) based on Jaccard (binary) distances calculated from the presence or absence of transposon insertions at TA sites (5,046 of 332,345 sites with >=3 reads in >=2 samples). Axes show the percentage of variance explained. Points are colored by week post-colonization and shaped by condition (Input, GF, SPF). **(E)** Relative abundance of transposon mutants mapping to *Pvu* genes across all samples after pooling mutants that map to the same gene. The 19 genes with the greatest relative abundance across all samples are separately colored, with other less abundant genes grouped in grey. **(F)** Transposon insertion map of the *Pvu2206-Pvu2208* locus showing TA insertion sites and mutant abundance in the input library and at weeks 1 and 4 following colonization of representative GF and SPF mice.

To accurately characterize the genetic determinants implicated in our Tn-seq screen, we re-sequenced *Pvu* CL09T03C04, integrating long- and short-read sequencing approaches. We generated a closed genome assembly consisting of a single contig. This updated assembly revealed minor differences in gene organization and genomic location compared with the previous annotation. Here and throughout the manuscript we use the updated gene designations; a complete cross-reference of previous and updated gene identifiers is provided in Supplementary Table S3.

To identify specific mutants contributing to the observed differences in mutant population structure, Tn insertion mutants mapping to the same gene were pooled and their relative abundance was quantified across all samples (Supplementary Table S3). We found that Tn insertion mutants in gene *Pvu2207* displayed a particularly striking environment-dependent phenotype (Figure 1E). In GF mice, *Pvu2207* insertion mutants became the dominant mutant population, accounting for a remarkable 25–75% of the recovered mutant library across all time points. In contrast, *Pvu2207* mutants were strongly depleted in SPF mice relative to the input library. Consistent with these abundance patterns, analysis of log2 fold-change values and p-values adjusted for multiple comparisons (<0.05) revealed that *Pvu2207* mutants were significantly enriched at weeks one and four in GF mice, while exhibiting significant depletion at weeks one and two in SPF mice (Figure 1C).

To prioritize candidate fitness determinants for downstream validation, we compared the whole-cell metabolomic profiles of several Tn mutants identified as differentially abundant in GF vs SPF mice from the screen using untargeted liquid chromatography with tandem mass spectrometry (LC-MS/MS) grown in rich media [BHIS: brain heart infusion (BHI) supplemented with hemin and vitamin K]. Transposon mutants were recovered from the arrayed library based on three criteria: a single transposon insertion in the genome, insertion near the N-terminus of the predicted coding sequence, and high abundance in either GF or SPF mice. Pairwise PERMANOVA identified significant differences in metabolite profiles between WT and mutants *Pvu255::Tn* and *Pvu2207::Tn*. Differences between WT and mutants *Pvu1650::Tn* and *Pvu32::Tn* were not statistically significant (Supplementary Figure S1F). The *Pvu255::Tn* and *Pvu2207::Tn* mutants exhibited significant differences in the abundance of metabolites belonging to the fatty acid amide, sphingolipid, phospholipid, and amino acid classes, based on FDR-adjusted statistical analysis (Supplementary Figure S1G and Supplementary Table S4). Among the mutants analyzed, the *Pvu2207::Tn* strain exhibited the greatest divergence from WT *Pvu* (Supplementary Figure S1F-G), supporting its selection for further characterization.

The hypothesis that insertions in *Pvu2207* were driving the *in vivo* phenotypes associated with this locus that were disparate between GF and SPF animals was further supported by the contributions of multiple unique Tn insertions in *Pvu2207* to the observed enrichment and depletion patterns (Supplementary Figure S1H and Figure 1F). Together, these results implicate *Pvu2207* as an environment-dependent fitness determinant whose contribution becomes apparent only within the context of a complex microbial community. These observations motivated further validation to determine whether *Pvu2207* directly contributes to *Pvu* fitness and to define the ecological and cellular processes underlying that phenotype.

### Genetic validation and transcriptional organization of the *Pvu2206–2208* locus

To determine whether the *Pvu2207::Tn* phenotype was specifically attributable to disruption of *Pvu2207*, we characterized the transcriptional consequences of the transposon insertion. Using the same *Pvu2207::Tn* mutant as in the untargeted metabolomics analysis above, RT-PCR analysis demonstrated that transcription of both *Pvu2207* and the downstream gene *Pvu2208* was disrupted, indicating that the *Pvu2207::Tn* transposon insertion generated a polar effect within the operon (Supplemental Figure S2A). To determine whether the upstream gene, *Pvu2206*, was transcribed along with *Pvu2207-Pvu2208* as a single operon, we performed RT-PCR using primer sets spanning the intergenic regions and gene boundaries. RT-PCR results indicate that transcription extends across the *Pvu2206–2208* region (Supplemental Figure S2A). The Prokaryote Promoter Prediction v2.0 (ProPr)^64^ tool predicted promoters immediately upstream of both *Pvu2206* and *Pvu2207* (Supplemental Figure S2B). In accord, RNA-seq of WT *Pvu* in rich growth medium, BHIS, as well as defined *Bacteroides* minimal medium (BMM)^29^ showed coverage of the entire *Pvu2206–2208* locus with marked increases in transcript levels for *Pvu2207–2208*. This suggests the promoter upstream of *Pvu2207* is stronger than the promoter upstream of *Pvu2206 in vitro* (Supplemental Figure S2B). Although *Pvu2205* is located immediately upstream of *Pvu2206*, RT-PCR analysis showed no detectable transcript spanning *Pvu2205* and *Pvu2206*, supporting our treatment of *Pvu2206–Pvu2208* as a distinct operon. Thus, *Pvu2205* was not investigated further. Together these data indicate that the locus comprises three co-transcribed genes, *Pvu2206*, *Pvu2207*, and *Pvu2208*.

To define the individual contributions of each gene within this locus, we generated clean, in-frame deletion strains using homologous recombination^48^. Four deletion strains were constructed: *Pvu*Δ*2206, Pvu*Δ*2207, Pvu*Δ*2208*, and *Pvu*Δ*2207-2208*. Each strain was validated by RT-PCR to confirm loss of transcription of the targeted genes (Supplementary Figure S2A). All deletion strains exhibited growth comparable to WT *Pvu* in both BHIS and BMM (Supplementary Figure S2C-D), suggesting that deletion of individual genes within this locus did not result in general *in vitro* growth defects.

### Functional predictions of the *Pvu2206–2208* locus provide insight into potential roles of the encoded proteins

Having identified *Pvu2207* as a potential microbiota-dependent fitness determinant, we renamed the genes within this locus “Determinant of Intestinal Fitness” (*dif*). To gain insight into the potential functions of proteins encoded by *Pvu2206-2208*, we generated functional predictions based on the amino acid sequences of each open reading frame using InterPro^43^, CDART^44^, Alphafold^38^, and eggNOG^45^. The protein product of Pvu2207 has a predicted N-terminal secretion signal sequence, but the remainder of the protein lacked a high-confidence predicted structure due to the N- and C- termini being disordered as identified by the intrinsic disorder prediction program, PONDR^39^ (Supplementary Figures S2E-G). However, when AlphaFold^38^ was combined with PONDR^39^ and IUPred3^40^, we found that Pvu2207, a 466 amino acid protein, contains a putative folded domain in the central portion (G63-Y187) that has partial homology to bacterial DUF3300 domain-containing proteins and animal vitellogenin II like-proteins as identified by FoldSeek^41^ (Supplementary Figure S2G-H). Given its unknown function and assignment to COG category S (function unknown), we designated *Pvu2207* as “*difS*”.

Given the predicted N-terminal secretion signal, we next examined the cellular localization of DifS using a FLAG-tagged strain, *PvuCL09difS-FLAG* (Supplementary Figure S2I-J). Dif*S*-FLAG was detected in the cell pellet but not the supernatant after lysis, suggesting that the protein is retained within the cell-associated fraction rather than secreted into the extracellular environment (Supplementary Figure S2K). This localization is consistent with a potential periplasmic or cell envelope-associated role for DifS, although its precise subcellular localization remains to be determined.

In contrast, functional predictions about the protein products of Pvu2206 and Pvu2208 were more informative. Pvu2208 is predicted with high confidence to fold into a β-barrel outer membrane protein containing a FadL domain, consistent with a potential role in transport of long-chain fatty acids or other hydrocarbons^65^ (Supplementary Figure S2F), so it was named “*difT*”, for “Transporter”. Pvu2206 was predicted with high confidence to belong to the HU/IHF superfamily of DNA-binding proteins (HU, histone-like protein; IHF, integration host factor) and to contain an HU-HIG domain placing it within a *Bacteroides* specific subclass of the greater HU/IHF superfamily^66^ (Supplementary Figure S2F). HU-HIG family members consist of a modified N-terminal HU domain linked to one or more of three distinct C-terminal domains: a winged helix-turn-helix (wHTH; H), an immunoglobulin-like fold (Ig; I), or a glycine-rich motif (G), which together give the clade its name^66^. Consistent with the first of these, InterPro^43^ assigned Pvu2206 to the winged helix-like DNA-binding domain superfamily (IPR036388), defining a wHTH domain at DifH’s C-terminus (Supplementary Figure S2F). Based on this predicted HU-HIG DNA-binding function, Pvu2206 was designated “*difH*”. Therefore, we hypothesized that multiple genes within this operon could contribute to *Pvu* fitness. We henceforth utilized the *difHST* nomenclature.

### Genomic organization and evolutionary conservation of the *dif* locus

To assess the conservation of this microbiota-dependent fitness locus across the *Bacteroidaceae*, we examined the presence and organization of the *difHST* locus in related species. We first determined the evolutionary relationship between *Pvu* and closely related species by comparing *Pvu* CL09T03C04 with 17 type strains within the *Bacteroidaceae* family using whole-genome sequences available through the Bacterial and Viral Bioinformatics Resource Center (BV-BRC^33^; Figure 2A). Phylogenetic analysis based on the 500 most conserved genes within these type strains identified two distinct *Phocaeicola* clades. *Pvu* CL09T03C04 and *Pvu* ATCC 8482 clustered with *P. dorei, P. sartorii,* and *P. massiliensis* (*Phocaeicola* Group 1), whereas *P. plebeius, P. barnesiae, P. coprophilus,* and *P. coprocol*a formed *Phocaeicola* Group 2. The remaining *Bacteroides* species formed a distinct *Bacteroides* Group (Figure 2A).

**Figure 2:**
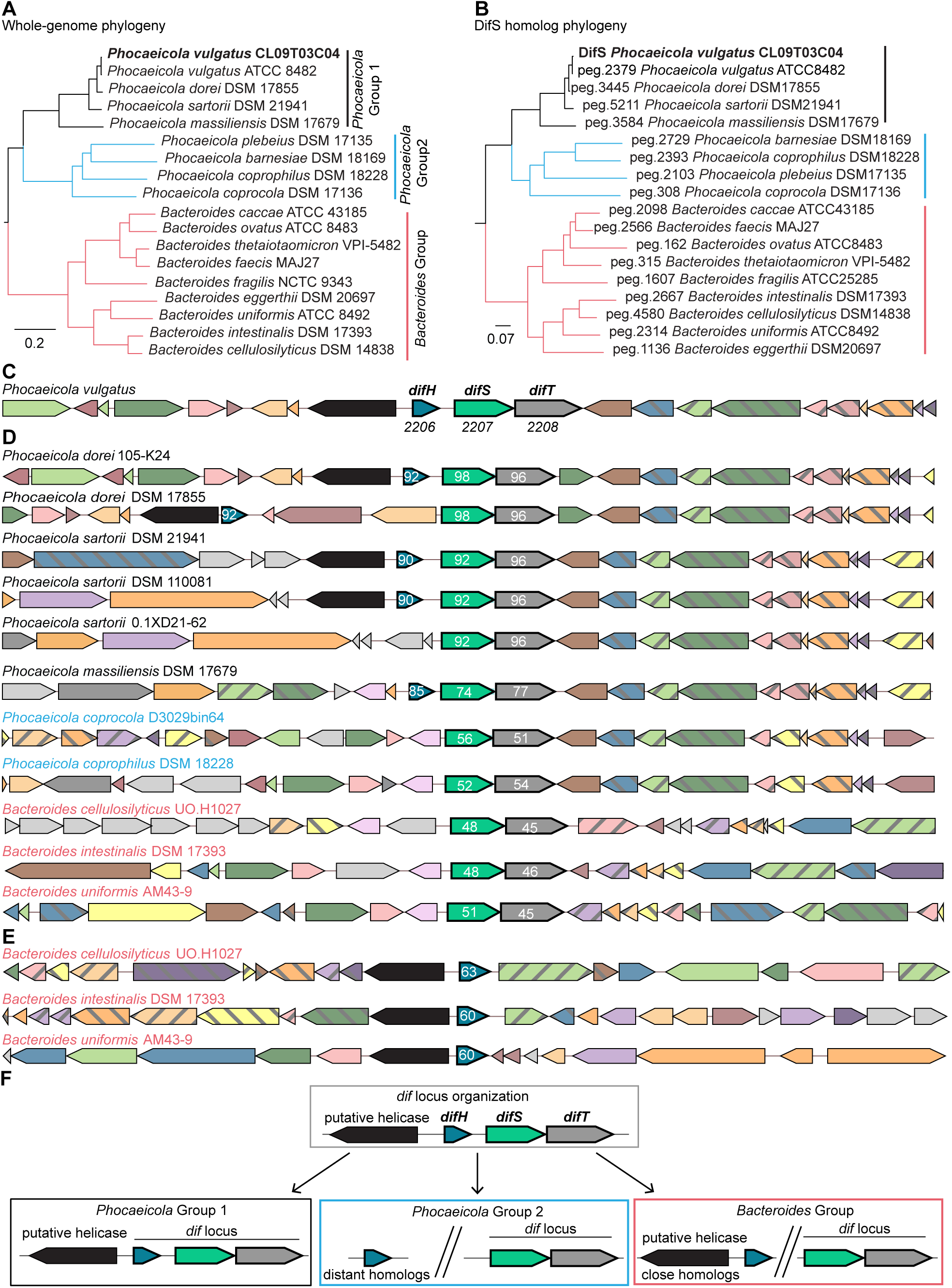
Conservation and genomic context of the *dif* locus in *Phocaeicola* and related *Bacteroidaceae.* **(A)** Whole-genome phylogenetic tree of select 17 type strains within the *Bacteroidaceae* family constructed using the 1,000 most conserved genes identified by BV-BRC. **(B)** Phylogenetic tree of DifS and homologous proteins from 17 *Bacteroidaceae* type strains identified by BLASTP (E-value ≤ 0.0001). **(C)** Genomic organization of the *dif* locus in *Pvu* CL09T03C04. **(D-E)** Conservation of the *dif* locus **(D)** and just *difH* **(E)** genomic neighborhood among representative *Phocaeicola* and closely related *Bacteroides* species. The numbers indicate percent similarity of that gene to each of the *dif* genes. **(F)** Graphic summary of the *dif* locus genomic neighborhood variations present in the *Phocaeicola* and *Bacteroides* species categorized into three distinct groups. Panels C-F were generated in Fast.genomics.

BLASTP identified DifS homologs in all 17 type strains, with their distribution broadly mirroring the *Bacteroidaceae* phylogeny (Figure 2B). We did not find clear homologs to DifS outside of the *Bacteroidota* phyla. Genomic neighborhood analysis using Fast.genomics^67^ revealed that *difS* and *difT* are adjacent in *Pvu* and all evaluated congeners. In contrast, *difH* is adjacent to the other two genes only in *Pvu* CL09T03C04 and other members of *Phocaeicola* Group 1 (Figure 2C–D). In some *P. dorei* and *P. sartori*i strains, all three genes occur within a contiguous locus, while others retain only the *difS-difT* pairing. Since BLASTP identified numerous DifH-related proteins across the evaluated genomes, we established sequence similarity criteria to distinguish close homologs from more distantly related homologs. Proteins sharing >40% amino acid similarity with DifH were operationally classified as “close homologs”, whereas proteins sharing 20–40% amino acid similarity were classified as “distant homologs”. Notably, *Phocaeicola* Group 2 contained only distant DifH homologs. In the *Bacteroides* Group, close *difH* homologs were present in the genome but not physically linked to their respective *difST* locus. In all species that possess a *difH* close homolog, we noted that *difH* is almost always adjacent to a putative helicase transcribed in the opposite orientation (Figure 2C–E). Together, these observations indicate that the *difS-difT* genomic arrangement is broadly conserved across *Bacteroidaceae*, whereas the *difH*-helicase region has a more restricted distribution within the *Phocaeicola* lineage (Figure 2F).

### Deletions within the *dif locus* reveal microbiota-dependent fitness defects during competition

We next sought to determine whether disruption of *Pvu2207* alone was sufficient to reproduce the fitness phenotype identified in the Tn-seq screen or whether additional genes within the operon also contributed, using single-gene and double-gene clean deletion mutants, respectively. We first tested the original *difS::Tn* compared to strains cleanly deleted for *difS*, *difH*, *difT,* and *difST* in GF and SPF mice. SPF mice that lacked endogenous *Pvu* were gavaged with each mutant strain independently, and all strains successfully engrafted and persisted at levels comparable to WT *Pvu* parental strain (Figure 3A). These results show that disruption of individual *dif* genes does not overtly impair intestinal colonization in the absence of direct competition with WT *Pvu*.

**Figure 3:**
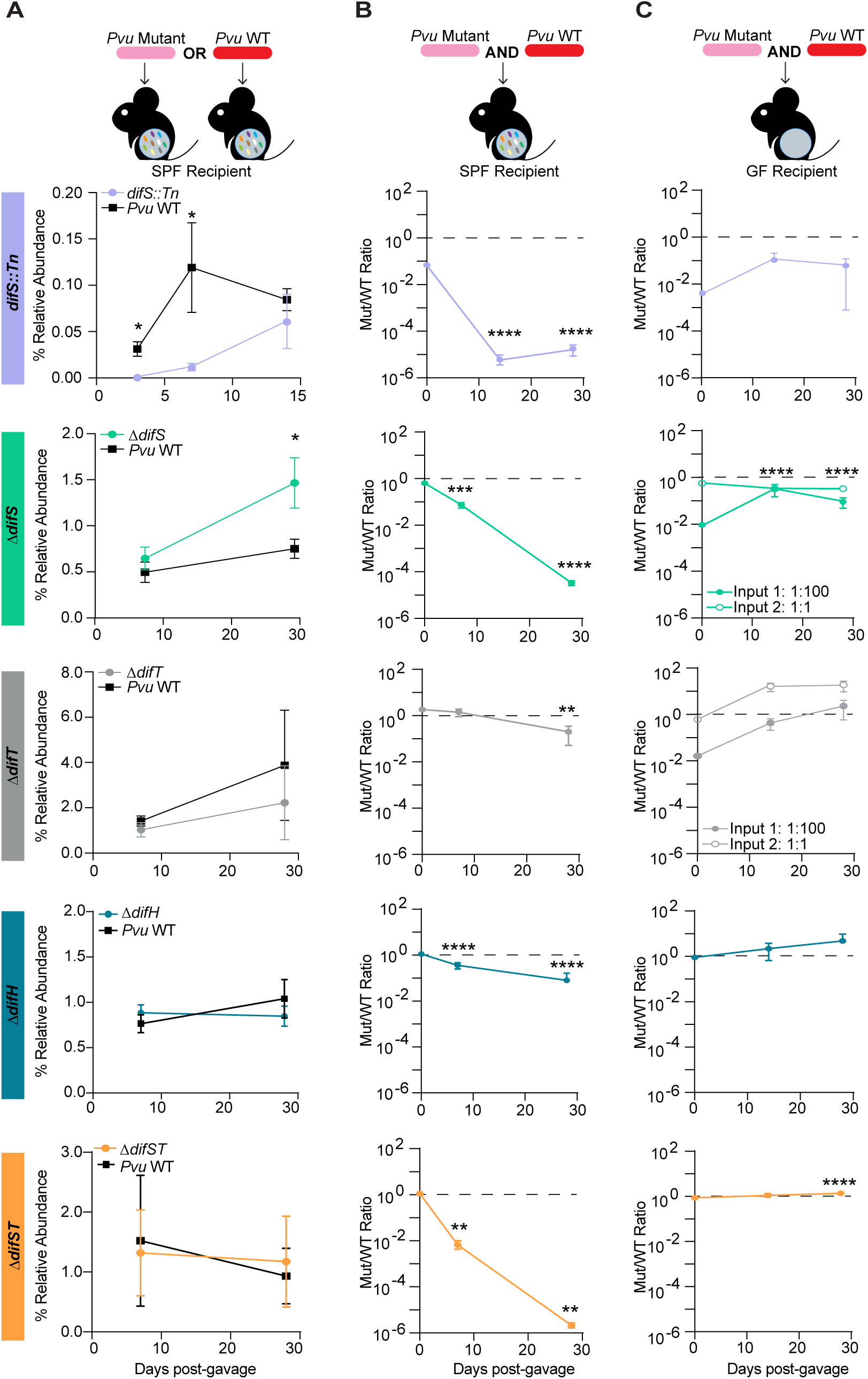
Deletion of genes in the *dif locus* reduces competitive fitness in SPF mice. **(A)** Relative abundance of WT *Pvu* and the indicated mutant strains following individual strain administration into SPF mice (n = 4-6). **(B-C)** Competitive fitness of the indicated mutant strains shown as a mutant/WT ratio following co-administration with WT *Pvu* into SPF (n = 4-12) **(B)** and GF mice (n = 3-6) **(C)**. Data are presented as mean ± SD. Statistical analysis was performed using an unpaired Mann-Whitney test for strain abundance at each time point **(A)** and paired Student’s t-tests comparing the Mut/WT ratio at each time point with the gavage ratio **(B–C)**. *p < 0.05, **p < 0.005, ***p < 0.001, ****p < 0.0001.

We next determined whether the clean deletion strains recapitulated the competitive fitness defects identified in the Tn-seq screen by competing each *dif* deletion directly against WT *Pvu* when given at a 1:1 ratio. Consistent with the Tn-seq results, all clean deletion strains exhibited reduced competitive fitness in SPF mice, although the magnitude of the defect varied among *dif* deletions (Figure 3B). The strongest fitness defects when competed against WT *Pvu* were observed for *difS::Tn,* Δ*difS*, and Δ*difST*. These data demonstrate that disruption of *difS* alone is sufficient to reproduce the phenotype identified in the Tn-seq screen. Notably, deletion of both *difS* and *difT* resulted in the greatest competitive disadvantage, suggesting that *difT* also contributes to the fitness phenotype associated with disruption of the *dif* locus.

In contrast, when the same *dif* deletion strains were competed against WT *Pvu* in GF mice, the competitive disadvantage of the *dif* deletion strains that was observed in SPF mice was abolished. The *dif* deletion strains maintained approximately 1:1 ratios with WT *Pvu* and, in some cases, increased in relative abundance when compared to the input ratio of WT to *dif* deletion (Figure 3C). These data are consistent with the increased abundance of *dif::Tn* observed in GF mice in the Tn-seq screen (Figure 1E). Additionally, when the *difS* and *difT* clean deletions were gavaged at a 1:100 ratio (mutant:WT) their relative levels increased up to or past a 1:1 ratio by 2 weeks post gavage (Figure 3C). Together, these findings demonstrate that disruption of *dif* genes does not impair intestinal colonization but does result in reduced fitness during microbiota-dependent intraspecies competition with WT *Pvu*.

### The competitive fitness defect of Δ*difS* is established in the colon within the first 24 hours following colonization

Having established that the *dif* locus influences the relative intraspecies competitive fitness within a complex microbial community, we sought to understand the timing and geography of that competitive interaction. We thus repeated the competition experiment between WT *Pvu* and Δ*difS* in SPF mice while collecting fecal samples at early time points during the first two weeks following oral gavage. A marked difference in strain abundance was evident at the first 24 hour timepoint after inoculation, with Δ*difS* exhibiting an approximate 30-fold reduction in relative abundance compared to WT *Pvu* as measured by qPCR (Figure 4A). These findings indicate that the competitive disadvantage of Δ*difS* manifests rapidly, within the first day following co-colonization with WT.

**Figure 4:**
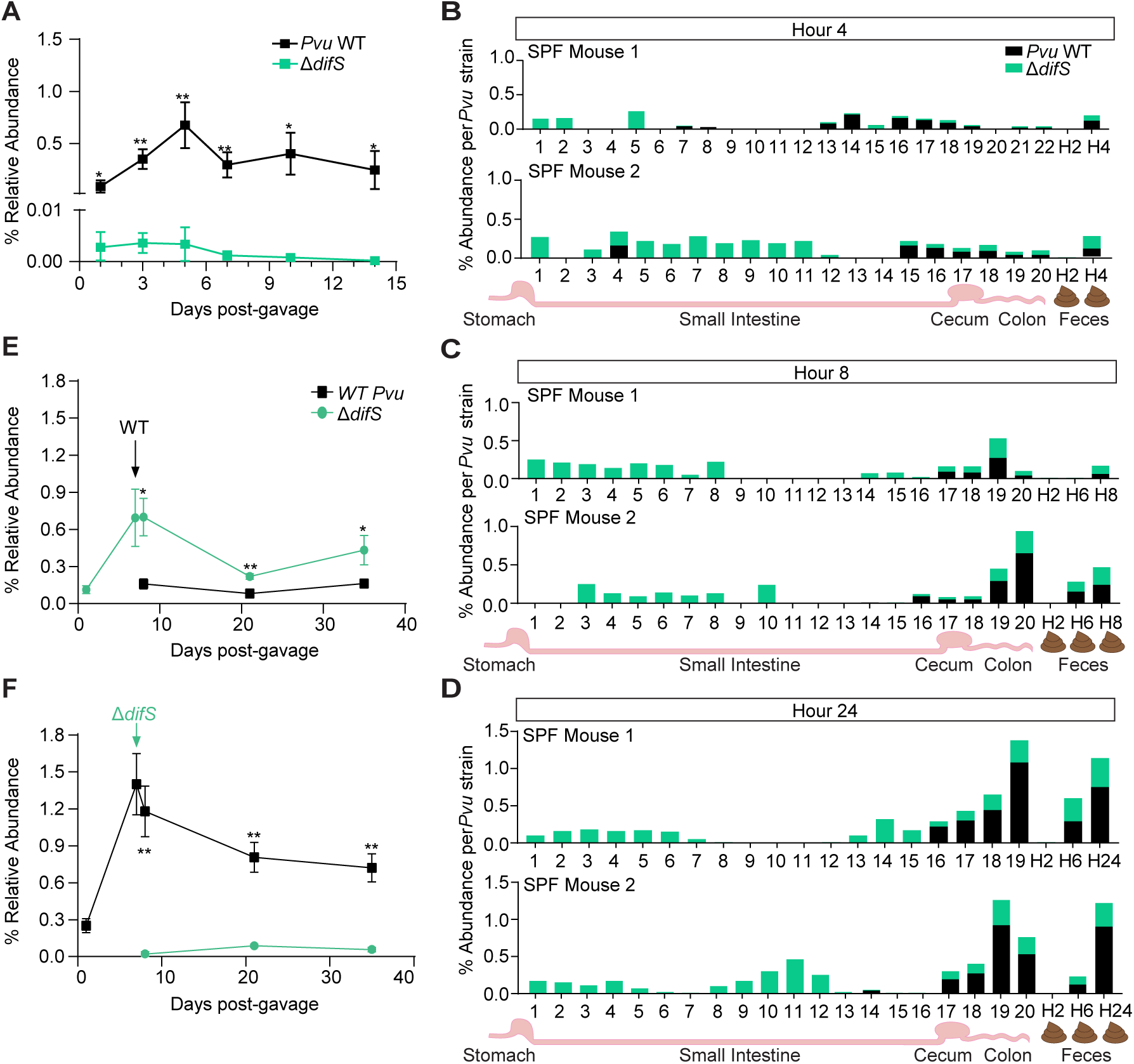
Rapid competition between WT and *ΔdifS* is associated with spatial separation and colonization order effects. **(A)** Relative abundance of WT *Pvu* and Δ*difS* during the first two weeks following co-administration at a 1:1 ratio into SPF mice (n = 5). **(B-D)** Spatial distribution of the relative abundance of WT *Pvu* and Δ*difS* along the intestinal tract at 4 hours (n=2) **(B),** 8 hours (n=2) **(C),** and 24 hours (n=2) **(D)** following co-administration into SPF mice. Relative abundance was determined from consecutive 2-cm intestinal sections extending from the duodenum to the rectum. **(E)** Relative abundance of WT *Pvu* and Δ*difS* following sequential administration into SPF mice (n = 5), with Δ*difS* administered one week before WT *Pvu* and **(F)** WT *Pvu* administered one week before Δ*difS* (n = 5). Data are presented as mean ± SD. Statistical analysis was performed using paired Student’s t-tests **(A, E-F)** *p < 0.05, **p < 0.005.

To resolve the early dynamics of this competition, SPF mice were again co-gavaged with WT *Pvu* and Δ*difS*, and fecal samples were collected every two hours for 24 hours post-gavage. In parallel, mice were sacrificed at 4, 8, and 24 hours post-gavage, and the entire intestinal tract was collected and sectioned into 2-cm segments from the duodenum to the rectum (Figure 4B-D). WT *Pvu* was detected exclusively within the ileum, cecum, and colon as early as 4 hours post-gavage. In contrast, Δ*difS* was detected across the intestinal tract, including the small intestine. This suggests that loss of *difS* alters the competitive fitness and intestinal distribution of *Pvu*, with the deletion strain unexpectedly showing greater representation in the small intestine. Despite this spatial segregation, the greatest differences in strain abundance occurred within the cecum and colon, where WT *Pvu* consistently maintained a higher abundance than Δ*difS* (Figure 4B-D) as expected. These findings demonstrate that differences in the intestinal distribution of Δ*difS* and WT *Pvu* are evident within 4 hours of gavage, with Δ*difS* showing greater localization to the small intestine and WT *Pvu* being more abundant in the colon and stool.

Since in these experiments both WT and Δ*difS* strains were introduced simultaneously, we next asked whether WT would similarly outcompete Δ*difS* if their colonization was temporally separated. We introduced either WT or Δ*difS* first into SPF mice, allowing them to establish for seven days before the competing strain was administered. Under these conditions, the strain colonizing first consistently maintained a substantially higher abundance than the strain introduced later, regardless of genotype (Figure 4E-F). Although the second strain remained detectable after gavage, it failed to reach levels comparable to the initial colonizer. Together, these findings indicate that *difS* contributes to intraspecies competition involved in *Pvu* niche establishment, as the reduced fitness associated with *difS* disruption is most pronounced during initial competitive colonization of the *Pvu* niche but becomes less apparent once an established *Pvu* population is present.

### Deletion of *difH* and *difS* lead to broad transcriptional changes in *Pvu*

Having identified the *dif* locus as a microbiota-dependent determinant of intraspecies fitness, we next sought to define the impacts of *dif* gene disruption on *Pvu* transcriptional programs. Since *difH* encodes a predicted DNA-binding protein and is located immediately upstream of *difS* and *difT,* we hypothesized that DifH may contribute to regulation of the locus or other genes involved in its fitness phenotype. We therefore performed RNA sequencing (RNA-Seq) on both Δ*difS* and Δ*difH* to characterize the transcriptional consequences of deleting each gene.

We found that the transcriptome of Δ*difS* clustered separately from WT *Pvu* and Δ*difH* under both minimal and rich media conditions, indicating that loss of *difS* resulted in broad changes to the transcriptional profile (Figure 5A). Although the media influenced the overall transcriptomic profiles, Δ*difS* was more distinct from both WT *Pvu* and Δ*difH* across both media conditions (Supplementary Figure S3A), suggesting that deletion of *difS* alters the transcriptional state regardless of nutrient availability.

**Figure 5:**
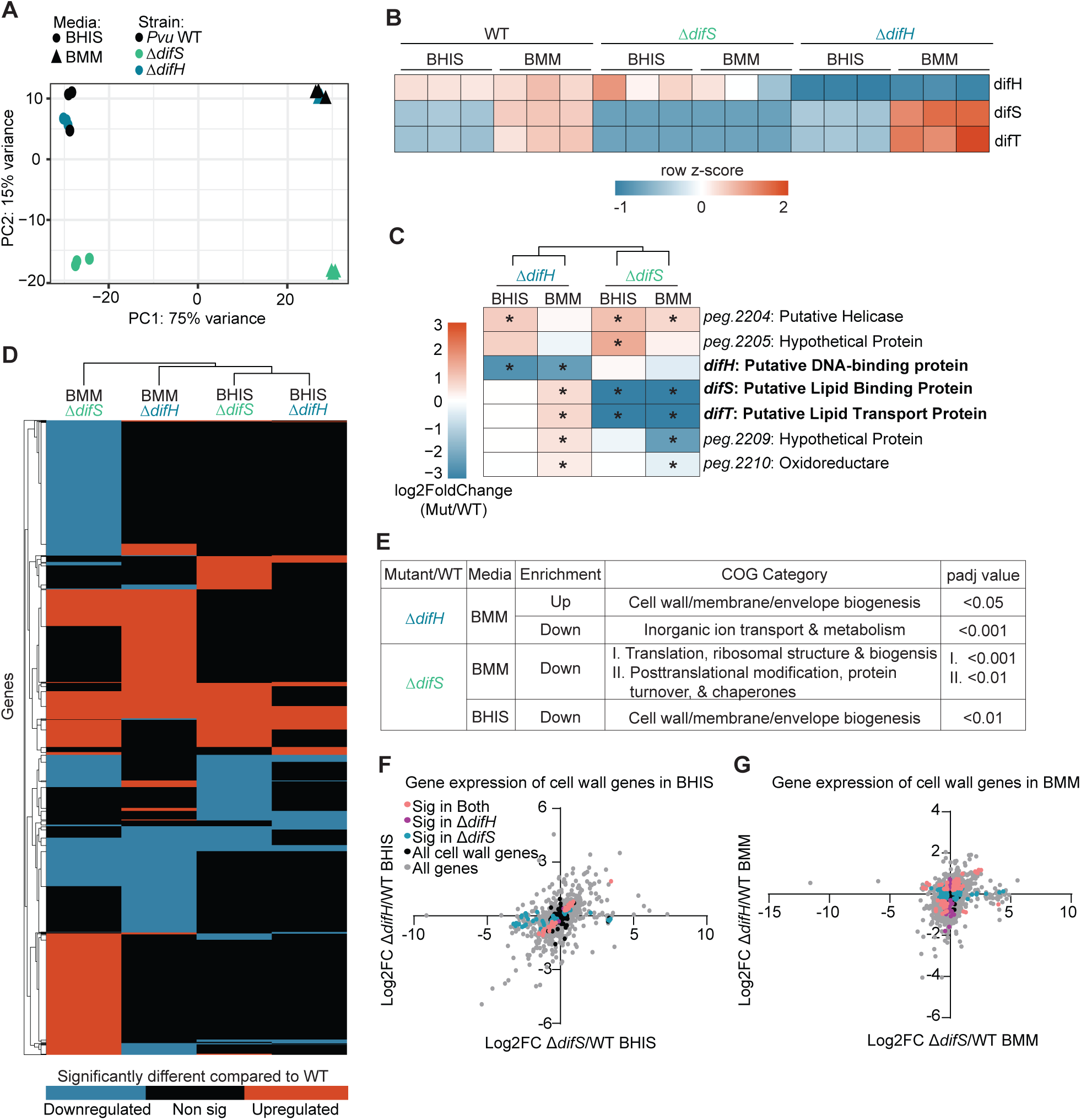
Deletion of *difS* and *difH* results in distinct transcriptional changes affecting operon regulation and cell envelope-associated functions. **(A)** PCoA of RNA-seq samples from WT *Pvu* (n = 3), Δ*difS* (n = 3), and Δ*difH* (n = 3) grown in BHIS and BMM. **(B)** Heatmap showing normalized transcript counts of *difH, difS*, and *difT* across biological replicates and growth conditions. **(C)** Heatmap showing log2FC values for genes within and surrounding the *dif* locus. **(D)** Heatmap showing genes with significant differential expression (adjusted p value < 0.05) in at least one deletion strain or growth condition. Log2FC values are shown, with blue indicating downregulation, orange indicating upregulation, and black indicating not significant. **(E)** COG enrichment analysis showing significantly altered functional categories in Δ*difS* and Δ*difH* compared with WT *Pvu* under BHIS and BMM conditions. **(F-G)** Scatterplots showing differential expression of all genes, including significantly and non-significantly differentially expressed genes, within the cell wall/membrane/envelope biogenesis COG category (COG M) in Δ*difS* (F) and Δ*difH* (G) compared with WT *Pvu* under BHIS and BMM conditions. Category enrichment was determined using a one-sided Fisher’s exact test. Significance: adjusted p values *, < 0.05; **, < 0.01; ***, < 0.001.

Next, we inspected the transcriptional regulation of the *dif* locus. In WT *Pvu*, *difH* was expressed at substantially lower levels than the downstream genes *difS* and *difT* and remained relatively unchanged between BHIS and BMM conditions. In contrast, *difS* and *difT* were expressed at higher levels in BMM than in BHIS (Figure 5B and Supplemental Figure S3B), indicating differential regulation of the three genes across growth conditions. Additionally, we saw that the deletion of *difS* resulted in significantly reduced expression of the downstream gene *difT* (Figure 5C). Upon further examination of our updated *Pvu* genome annotation, we noticed that the deletion boundaries used to originally generate Δ*difS* overlapped a region of 129 base pairs that was predicted to contain part of the internal promoter upstream of *difS.* Thus the reduced expression of *difT* in Δ*difS* may be due to reduced transcriptional activity from the altered shared *difS* promoter rather than being due to loss of *difS* gene function. However, the positive PCR product for *difT* seen using cDNA from Δ*difS* as template (Supplemental Figure S2A) indicates that *difT* is still expressed in the Δ*difS* strain. Interestingly, deletion of *difH* resulted in significantly increased expression of both *difS* and *difT* under BMM conditions, suggesting a potential negative regulatory effect of *difH* on expression of the downstream genes *difST* (Figure 5C). Together, these observations suggest that while *difS* and *difT* exhibit increased expression under nutrient-limited conditions, they also may be negatively regulated by *difH*.

We next compared the broader transcriptional changes associated with deletion of *difS* and *difH* relative to WT *Pvu* (Supplementary Table S5). Both of these strains exhibited extensive transcriptional changes, with hundreds of genes significantly upregulated or downregulated (Supplemental Figure S3C-D). Differential gene expression patterns differed between Δ*difS* and Δ*difH* and across growth conditions (Figure 5D, Supplementary Figure S3C, Supplementary Table S5). Several genes exhibited opposing expression patterns between the mutants, suggesting distinct and potentially opposing roles for DifS and DifH in transcriptional regulation (Figure 5D; Supplemental Figure S3C; Supplementary Table S5). Furthermore, the transcriptional response of Δ*difH* varied more substantially between rich and minimal media than that of Δ*difS*, suggesting that *difH* has a more media-dependent regulatory effect (Supplementary Figure S3E-F).

To better understand the biological processes influenced by *difH* and *difS*, we performed COG enrichment analysis^68,69^ on differentially expressed genes associated with each deletion strain. First, each strain was compared across the two media conditions, BMM versus BHIS, and saw that when the strains are grown in nutrient limiting conditions, there is a downregulation of categories associated with metabolism and transport (Supplementary Figures S3G-I). Conversely, expression of genes associated with transcription was increased following deletion of *difH* in BMM (Figure S3H), consistent with a potential role for *difH* as a negative regulator of transcription. In contrast, deletion of *difS* resulted in increased expression of genes associated with inorganic ion transport and metabolism as well as replication, recombination, and repair (Supplementary Figure S3I).

Comparing each deletion strain against WT *Pvu*, we found that the COG category associated with cell wall, membrane, and envelope biogenesis (COG Category M; Supplementary Table S5) was significantly altered in both mutants, but in opposite directions, with enrichment in Δ*difH* and depletion in Δ*difS* (Figure 5E-G and Supplementary Figure S3J-K). These differences were more pronounced in BMM, where Δ*difH* showed broader upregulation of genes involved in outer membrane proteins, efflux and transport, glycan biosynthesis, and other cell envelope functions, whereas many of these genes were downregulated in Δ*difS* (Figure 5F-G; Supplementary Table S5). Fourteen genes showed negatively correlated expression patterns between the two deletion strains relative to WT, with most involved in glycan biosynthesis and capsule-associated functions (Supplementary Table S5). Given that DifH is a DNA-binding protein in the same locus as *difS-difT*, these opposing transcriptional responses suggest that DifH and DifS influence overlapping cell envelope functions with distinct regulatory effects. Together, these findings indicate that the *dif* locus contributes to transcriptional regulation of cell envelope homeostasis, with the effects of *difH* and *difS* deletion differing in both magnitude and direction.

### Mutant strains exhibit limited changes in cellular lipid and metabolite profiles

Our observations above all indicated that the genes in the *dif* locus are important for regulating cell wall- and membrane-associated genes. We next asked whether these gene expression shifts in the *dif* deletion strains relative to WT *Pvu* were accompanied by changes in the production of specific cellular lipids or metabolites. We performed untargeted lipidomics and metabolomics on *Pvu* cell pellets collected following *in vitro* growth. Since the transcriptional data indicated that differences between *dif* deletion strains and WT strains were most apparent under BMM conditions, we selected BMM to conduct metabolomics and lipidomics experiments. The choice of this defined media also simplified mass spectrometry analysis, since this defined growth medium contained no exogenous lipids, making potential detection of changes in cellular lipids associated with the *dif* deletions more straightforward.

PCA of the overall lipidomic and metabolomic profiles showed that the *dif* deletion strains all clustered with WT *Pvu* (Supplementary Figure S4A–B). Pairwise PERMANOVA of whole-cell lipidomic profiles revealed no significant differences between WT *Pvu* and the *ΔdifS*, *ΔdifT*, or *ΔdifST* mutants after multiple-testing correction (BH-adjusted *p* > 0.05). Among these mutants, *ΔdifT* exhibited the greatest separation from WT in lipidomic profiles, although this difference was not statistically significant (F = 1.76, R² = 0.180, BH-adjusted *p* = 0.225). Similarly, pairwise PERMANOVA of whole-cell metabolomic profiles identified a significant difference between WT *Pvu* and the *ΔdifST* mutant (F = 9.44, R² = 0.486, BH-adjusted *p* = 0.010), whereas the *ΔdifH*, *ΔdifS*, and *ΔdifT* mutants did not differ significantly from WT after multiple-testing correction (BH-adjusted *p* > 0.05). Together, these findings indicate that deletion of the *dif* locus genes did not substantially alter the overarching lipidomic or metabolomic profiles of *Pvu*.

Lipid-specific methods did not reveal any specific lipid features that were statistically significantly different between strains (Supplementary Table S5). Consistent with this, a comparison of broader categories of membrane or lipid classes did not indicate substantial differences in membrane composition between the *dif* deletion strains and WT (Supplementary Figures S4C-E). Although *ΔdifT* appears to be the most different from WT with respect to the overall membrane composition of cell pellets (Supplementary Figure S4C), these differences were not statistically significant. Global metabolite profiling in BMM did identify several significant differences across multiple metabolite classes, including lipids, amino acids, nucleosides and nucleotides, and organic acids (Supplementary Figure S4F and Supplementary Table S5). Notably, the significant lipid differences in *ΔdifST* included a reduction of several lyso-PE (1-Palmitoyl-2-hydroxy-sn-glycero-3-phosphoethanolamine), lyso-PC (1-Hexadecyl-sn-glycero-3-phosphocholine), lyso-PS (PS(18:0/0:0)), and monoacylglycerol (Glycerol 1-myristate / monolaurin) species (Supplementary Figure S4F), consistent with potential alterations in phospholipid turnover or synthesis.

Overall, these data suggest that while there are shifts in some metabolic features between WT and *dif* deletion strain, the changes are not substantial enough to cause large-scale lipid remodeling under the *in vitro* conditions tested. One important caveat to these conclusions is that our analysis focused exclusively on features that are annotatable based on currently available databases. Our data do reveal unannotated features that are significantly different between the WT and the *dif* deletion strains. In the absence of validated identifications for these features, we were unable to make conclusions about their meaning or significance at this time.

### Deletion of *difS* increases susceptibility to cell envelope targeting agents

Having identified transcriptional changes associated with cell envelope homeostasis, we next sought to test directly whether disruption of the *dif* locus resulted in measurable membrane-associated phenotypes. To test this, WT *Pvu* and the *dif* deletion strains were challenged with a panel of cell envelope targeting antibiotics and physiologically relevant bile acids to compare growth responses under BHIS and BMM conditions.

We first examined susceptibility to polymyxin B, carbenicillin, meropenem, and fosfomycin, which target components of the cell envelope (Figure 6A–D). Across assays for the first three antibiotics in BHIS, Δ*difS* and Δ*difST* consistently exhibited the greatest growth defects, whereas Δ*difH* behaved similarly to WT *Pvu.* Δ*difT* displayed an intermediate phenotype, suggesting that *difS* is the primary determinant of this envelope-associated phenotype, with *difT* also contributing (Figure 6A-C). All strains were resistant to fosfomycin, suggesting that the *dif* genes do not contribute to the initial step of peptidoglycan biosynthesis (Figure 6D). Interestingly, these differences were largely absent when strains were grown in BMM, as all strains ultimately recovered following antibiotic exposure (Figure 6A-D). Evaluation of multiple antibiotic concentrations demonstrated that Δ*difS* and Δ*difST* consistently exhibited greater sensitivity than the other strains across a range of doses (Supplementary Figure S5A-D).

**Figure 6:**
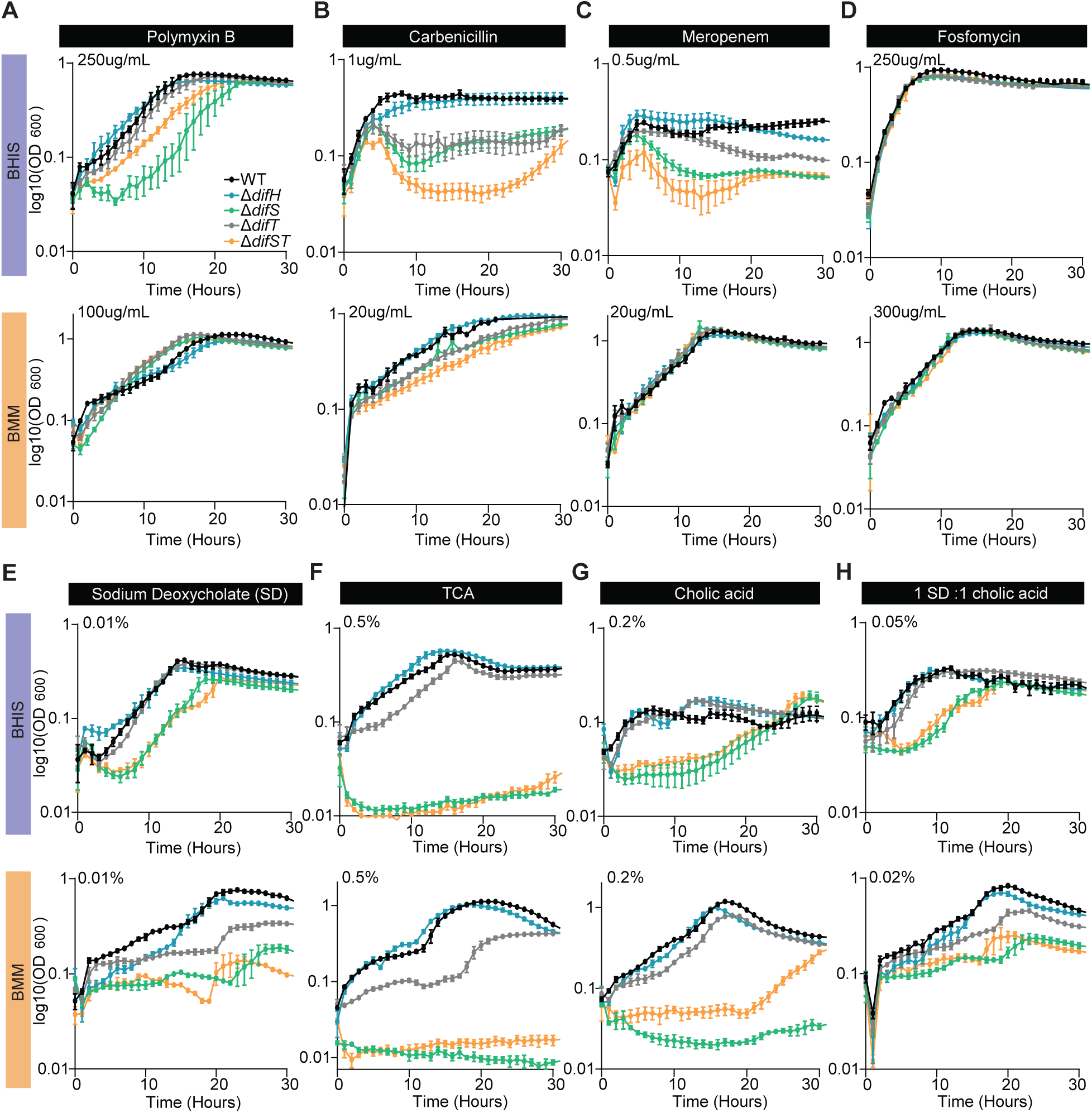
Deletion of *difS* increases sensitivity to membrane-targeting compounds and intestinal stressors. **(A-D)** Growth curves of WT *Pvu* and *dif* deletion strains (n = 3) in the presence of membrane- or cell envelope-targeting antibiotics: 250ug/mL of polymyxin B **(A),** 1ug/mL of carbenicillin **(B),** 0.5ug/mL of meropenem **(C),** and 250ug/mL of fosfomycin **(D)** tested in BHIS and BMM. **(E-F)** Growth curves for WT *Pvu* and *dif* deletion strains (n = 3) for 4 different physiologically relevant salts: 0.01% sodium deoxycholate **(E),** 0.5% TCA **(F),** 0.2% cholic acid **(G),** and 0.05% 1:1 sodium deoxycholate:cholic acid **(H)** tested in BMM and BHIS.

Due to their ability to disrupt bacterial membranes^70,71^, we next examined susceptibility to bile acids commonly encountered by intestinal bacteria, including sodium deoxycholate, taurocholic acid (TCA), cholic acid (CA), and a 1:1 mixture of sodium deoxycholate and cholic acid (Figure 6E-H). In contrast to the antibiotic assays, Δ*difS* and Δ*difST* displayed pronounced growth defects in both BHIS and BMM, whereas WT, Δ*difH*, and Δ*difT Pvu* exhibited comparable growth. As observed for the antibiotic assays, dose-response experiments consistently demonstrated increased sensitivity of Δ*difS* and Δ*difST* across multiple concentrations (Supplementary Figure S5E-H). Together, these findings demonstrate that disruption of *difS* and *difST* increases susceptibility to a broad range of envelope-targeting antibiotics and physiologically relevant bile compounds, providing functional evidence that this operon contributes to maintenance of membrane integrity under environmental stress.

### DifH defines a divergent lineage of IHF-like proteins in *Bacteroidota*

RNA-seq analysis revealed that deletion of *difH* resulted in increased expression of *difS* and *difT*, suggesting that DifH may repress transcription of its downstream genes (Figure 5C). Since these genes contribute to microbiota-dependent intraspecies fitness, these findings raised the possibility that DifH influences the *dif* locus through transcriptional regulation of *difS* and *difT*. This observation prompted us to place DifH within the broader HU/IHF family. We therefore performed a phylogenetic analysis of DifH alongside IHFα and IHFβ proteins and the broader group of HU/IHF proteins previously described^66^, and all close DifH homologs as identified by BLASTP (Supplementary Figure S6A).

Although DifH is part of the HU/IHF superfamily, it did not cluster with the canonical IHFα and IHFβ proteins from *E. coli* (Supplementary Figure S6A). Instead, DifH fell into a distinct subclade of homologs from *Phocaeicola* and *Bacteroides* species. Notably, sequence relatedness among these homologs did not strictly correspond to genus-level taxonomy. The closest DifH homologs were identified in *P. dorei, P. sartorii,* and *P. massiliensis* (*Phocaeicola* Group 1; Fig. 2A). Related proteins from other *Phocaeicola* species, including *P. barnesiae, P. coprophilus, P. plebeius*, and *P. coprocola* (*Phocaeicola* Group 2; Fig, 2A), were substantially less similar to *Pvu* DifH (distant homologs). Moreover, the proteins most closely related to DifH also contain the HU-HIG domain and the winged helix-like domain (wHTH; InterPro: IPR036388; G3DSA:1.10.10.10) (Supplementary Figure S6B)^46,47^. Thus DifH represents a distinct wHTH clade of HU/IHF proteins within the *Bacteroidota* with a distribution that does not strictly follow species relatedness (Figure 2F).

### DifH binds near genes involved in metabolism, envelope biogenesis, and cellular defense

We next sought to directly define the genomic targets of DifH using chromatin immunoprecipitation sequencing (ChIP-seq). We used homologous recombination to insert an in-frame HA epitope tag at the C-terminus of *difH in situ* and under control of its native promoter^48^. Expression of the tagged protein was confirmed by western blot (Supplementary Figure S7A) and RT-PCR confirmed that insertion of the HA tag did not alter transcription of the locus (Supplementary Figure S7B). Further, the tagged strain grew comparably to WT *Pvu* in both BHIS and BMM (Supplementary Figure S7C-D), and *PvuCL09difH-HA* exhibited fitness comparable to WT *Pvu* in competitive colonization experiments in SPF mice (Supplementary Figure S7E). Together, these results suggest that the HA-tagged strain retained WT-like function and was suitable for immunoprecipitation.

Using the validated *PvuCL09difH-HA* strain, we performed ChIP-seq following growth in BHIS and BMM using a modified version of an established protocol^50^. Pooled biological replicates were analyzed with Model-based Analysis of ChIP-Seq 2 (MACS2)^56,57^ to identify genomic regions enriched for DifH binding relative to pooled WT samples. MACS2 identified 1,319 candidate binding sites in BHIS and 27 in BMM (q-value = 0.001) (Figure 7A-B; Supplementary Figure S8A-D; Supplementary Table S7). After controlling for background signal reduction (Supplementary Figure S8A-B), and in combination with peak analysis using Multiple Em for Motif Elicitation (MEME)^61,62^, we identified that only 14 of the BHIS peaks and three of the BMM peaks contained a recognized motif (Figure 7A-B, Supplemental Figure S8C-D, and Supplementary Table S7). Notably, the three strongest DifH binding sites identified in BHIS corresponded to the same sites detected in BMM, including one located immediately upstream of *difH* at the start of the *dif* operon, indicating that DifH consistently occupies these loci across growth conditions and binds its own promoter region (Figure 7C-E). The remaining highest-confidence peaks were located proximal to genes involved in carbohydrate transport and metabolism, lipid transport and metabolism, cell wall/membrane/envelope biogenesis, amino acid transport and metabolism, and nucleotide transport and metabolism, as well as genes associated with defense mechanisms (type I restriction-modification and CRISPR-Cas systems) and mobile genetic elements (Supplementary Table S7). Notably, one of the high-confidence DifH binding peaks occurred in the region immediately upstream from the gene encoding one of the close DifH homologs, *peg.598.* Further inspection of the genomic loci encoding other close DifH homologs in *Pvu* CL09 (Supplementary Figure S6A) revealed that only *peg.598* is arranged in a similar configuration as *difH* (Figure 2C) with an adjacent predicted helicase gene (*peg.596*) transcribed in the opposite direction. We also noticed that the locations of those strongest DifH binding peaks on the *Pvu* chromosome were often in close proximity (e.g., *peg.1110* and *peg.1125-6*, *peg.587* and *peg.623-4*; Supplementary Table S7), raising the possibility of physical interaction and DNA looping between adjacent DifH binding sites. Together, these findings identified a limited set of conserved genomic loci directly bound by DifH, including its own promoter.

**Figure 7:**
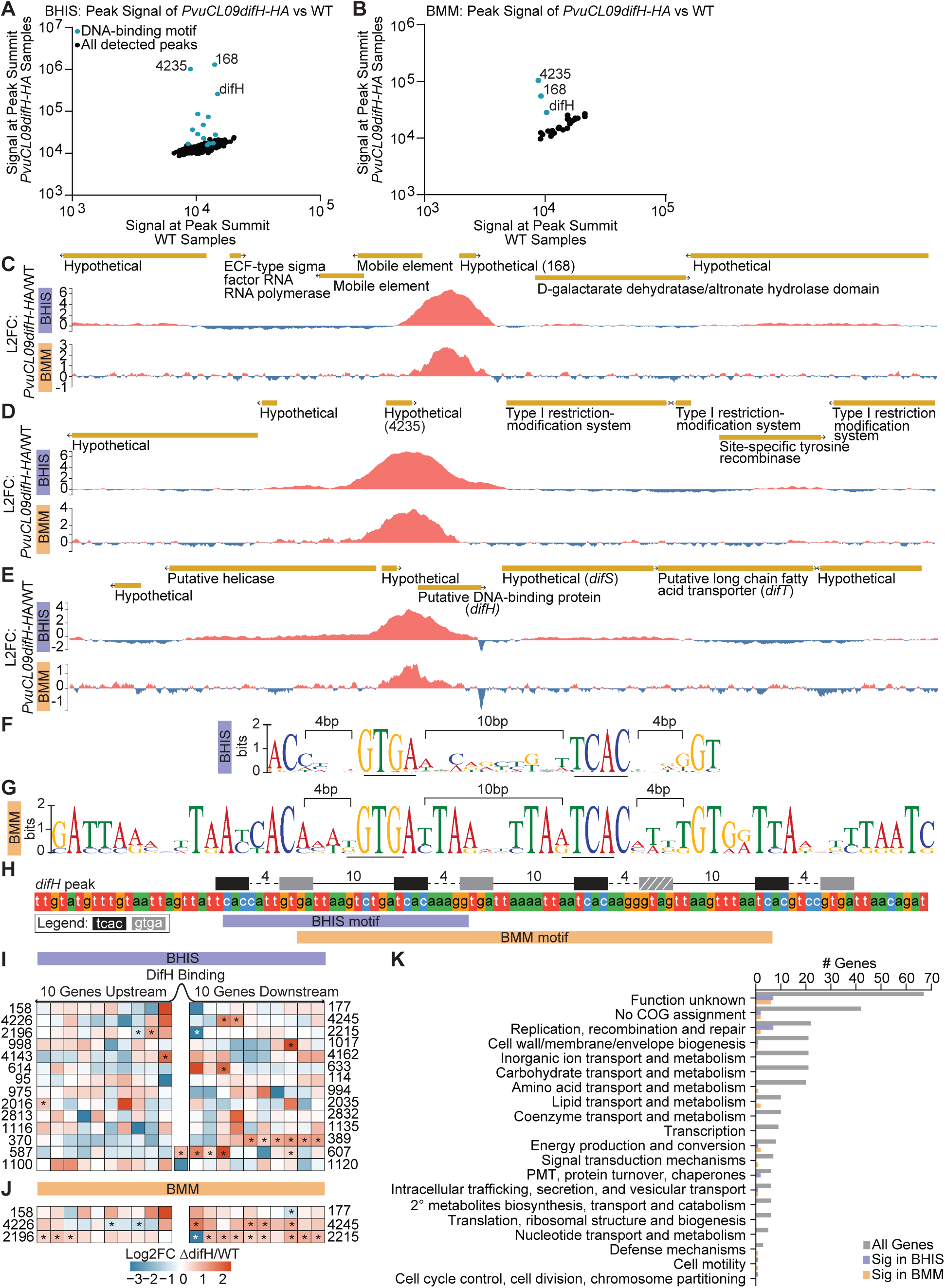
DifH binds upstream of the *dif* operon and additional loci associated with cellular defense functions. **(A-B)** Peak summit enrichment for all ChIP-seq peaks identified by MACS2 (q < 0.001) from *difH-HA* (n = 4) and WT *Pvu* (n = 4) grown in BHIS **(A)** and BMM **(B).** Peaks containing the MEME-identified DNA-binding motif are highlighted in blue. **(C-E)** ChIP-seq genome browser tracks of the three most enriched DifH binding sites in BHIS and BMM. Tracks show log2 fold enrichment of *difH-HA* relative to WT *Pvu*. **(F)** Sequence logo of the 30-bp DNA-binding motif identified by MEME from the top 75 BHIS ChIP-seq peaks (E-value = 4.5 × 10^-16^). **(G)** Sequence logo of the 58-bp DNA-binding motif identified by MEME from all 27 BMM ChIP-seq peaks (E-value = 2.4 × 10^5^). **(H)** DNA sequence surrounding the identified DifH-binding motif upstream of the *difH* gene in both BHIS and BMM. The motif consists of two highly conserved sequences, GTGA (gray boxes) and TCAC (black boxes), separated by 10 bp, as indicated by solid lines. Successive motif units are spaced 4 bp apart, as indicated by dashed lines. Boxes with white lines indicate instances in which either conserved sequence contains a mismatch or inversion. **(I-J)** Gene neighborhoods surrounding DifH ChIP-seq peaks containing the DNA-binding motif, overlaid with RNA-seq log2 fold-change values for Δ*difH* relative to WT *Pvu* in BHIS **(I)** and BMM **(J).** For each peak, the 10 genes upstream and downstream are shown. DifH peaks are located within intergenic regions for all but the two bottom loci, in which the motif-containing peaks fall within coding sequences. Asterisks denote significantly differentially expressed genes determined using a one-sided Fisher’s exact test (adjusted p < 0.05). White asterisks denote significant expression of DifH. **(K)** COG categories of the 10 up- and downstream genes of each peak. The genes that are significantly differential in Δ*difH* relative to WT *Pvu* in BHIS (purple) and BMM (yellow) are shown.

In BHIS, motif analysis identified a 30-bp palindromic sequence characterized by highly conserved GTGA and TCAC residues flanking a more variable 10 base pair central region (Figure 7F and Supplementary Figure S8E). Although only three out of the 27 binding sites were identified to contain a motif in BMM, the associated 58-bp motif retained the same conserved GTGA/TCAC regions with the same 10 base pair gap between them despite greater overall variability (Figure 7G and Supplementary Figure S8F), suggesting that DifH recognizes a common DNA sequence under both growth conditions. Appraisal of the position weight matrices for the 30-bp and 58-bp motifs (Figure 7F-G) suggested the possibility of an even larger motif with a repetitive structure under those peaks. We therefore manually inspected the DNA sequence content within each of the DifH binding peaks which contain those motifs. This revealed a larger structure consisting of repeating units of the identified GTGA(10-bp)TCAC block separated by 4-bp gaps at the *difH* promoter (Figure 7H) and the other sites of DifH occupancy (Supplementary Figure S9). With decreasing DifH occupancy, we observed a trend towards increased degeneracy of the GTGA/TCAC motif and reduced numbers of those repeating units (Supplementary Figure S9). These results identify a DifH DNA-binding motif and suggest that DNA occupancy may be enhanced by strictly spaced clustering of multiple motifs. Given the presence of DifH homologs and other HU/IHF proteins in additional *Phocaeicola* and *Bacteroides* species, we asked whether the DifH DNA-binding motif is conserved across these genomes and how frequently it occurs. The 30-bp DifH DNA-binding motif (Figure 7F) was detected in *Pvu* CL09T03C04 and all 17 additional type strains examined, with frequencies ranging from 61 to 82 sites per megabase with *Pvu* specifically having 78.5 motifs per megabase (Supplementary Figure S8G). The frequent occurrence of the DifH DNA-binding motif across the genomes of all strains examined indicates that sequences resembling the DifH binding motif are broadly conserved across these 17 species. However, individual motif occurrence alone does not establish DifH occupancy or regulatory activity, as our ChIP-seq revealed that only a small specific subset of those motifs are significantly occupied by DifH under *in vitro* conditions (Figure7A-B).

We then integrated the ChIP-seq and RNA-seq datasets to determine whether direct DifH binding correlated with transcriptional changes observed following deletion of *difH.* For each binding site containing the DNA motif in BHIS, we examined differential gene expression within an arbitrary 10-gene neighborhood around each binding site (Figure 7I-J and Supplementary Table S6). Significant transcriptional changes in Δ*difH* were observed in the neighborhoods surrounding some, but not all, binding sites. Moreover, neighboring genes frequently displayed mixed transcriptional responses, with some genes exhibiting increased expression while others were reduced (Figure 7H-I). As noted above (Figure 5C), deletion of *difH* led to increased expression of *difST* and several adjacent genes in BMM but not BHIS (Figure 7I-J). Notably, one of the largest effects of *difH* deletion was at the locus of its close homolog *peg.598* (Supplementary Figure S6A), where transcripts for *peg.597-600* were significantly elevated in Δ*difH* mutants (Figure 7I and Supplementary Table S7). These findings suggest that DifH binding is associated with transcriptional regulation at a subset of genomic loci rather than uniformly activating or repressing neighboring genes.

Finally, we next examined the COG designations of these up- and downstream genes, as well as genes identified as significantly differentially expressed in the Δ*difH* vs WT RNA-seq comparison from both BHIS and BMM (Figure 7K). The majority of genes either lacked a COG assignment or were classified as Function Unknown (Category S; Supplementary Table S4), consistent with the incomplete annotation of the *Pvu* genome. Among the assigned COG categories, the largest numbers of genes fell within replication, recombination, and repair (Category L) and cell wall/membrane/envelope biogenesis (Category M), suggesting that DifH may regulate genes involved in cellular defense and cell envelope functions (Supplementary Table S6). Together, these findings indicate that although DifH directly binds a limited number of genomic loci, deletion of *difH* is associated with broader transcriptional changes, consistent with a role for DifH in regulating cellular envelope functions and defense-associated processes through both direct and indirect effects.

## DISCUSSION

A comprehensive understanding of the mechanisms governing gut microbial ecology requires insight into the fitness landscapes encountered by its major bacterial constituents. Our findings expand this framework by identifying a previously uncharacterized locus encoding genes associated with intestinal fitness and cell envelope resilience. This *dif* locus is conserved among related *Phocaeicola* species and its disruption alters the expression of genes involved in cell envelope, membrane, and defense-associated functions. These changes are accompanied by defects in fitness during intraspecies competition in the mouse gut. Together, these findings establish the *dif* locus as a determinant of *Pvu* fitness and implicate cell envelope homeostasis in the ability of *Pvu* to persist against conspecific competitors in the gut.

Several limitations should be considered when interpreting our findings. First, interpretation of the Tn-seq and RNA-seq data is constrained by the incomplete annotation of the *Pvu* genome, with only approximately 30–40% of genes currently annotated. This limits our ability to assign functions to uncharacterized genes and fully interpret the pathways associated with the observed phenotypes. Second, although our *in vitro* and *in vivo* analyses collectively implicate the *dif* locus in *Pvu* fitness, the extent to which the phenotypes observed under laboratory growth conditions reflect those occurring in the gut remains unclear. Further studies are needed to establish how these phenotypes relate across biological contexts. Third, the murine gut differs from the human gut in bile acid composition, diet, transit time, and resident microbiota, which is particularly relevant given the bile salt sensitivity phenotype central to this work. Each cage housed a distinct strain or strain combination, so cage and strain were aligned in this design. Potential cage effects were not assessed separately from strain effects. In addition, because mice are coprophagic, some exchange of microbiota among cage mates may occur. Finally, although transcriptomic and ChIP-seq analyses provide insights into potential regulatory relationships involving DifH and cell envelope-associated genes, the precise molecular mechanisms linking the *dif* locus to fitness during intestinal colonization remain to be resolved.

Our *in vivo* results suggest that the *dif* locus is not required for *Pvu* to access or establish itself within the mouse intestine but instead becomes important when *Pvu* must compete for a niche already occupied, or actively being occupied, by another *Pvu* strain. The ability of *dif* deletion strains to colonize the SPF intestine in the absence of WT *Pvu* may reflect the availability of an unoccupied *Pvu*-specific niche, allowing even these competitively disadvantaged *dif* deletion strains to establish when no other competing *Pvu* strains are present. This interpretation is consistent with the unusual ability of *Pvu* to invade an established microbiota following a single introduction^8,17^. It also suggests that *Pvu* has access to a substantial permissive niche within the gut microbiome that it can readily occupy in the absence of a pre-existing resident *Pvu* population. Notably, loss of *difS* also altered the spatial distribution of *Pvu* within the intestine, with WT *Pvu* detected primarily in the ileum, cecum, and colon, whereas the colonization of Δ*difS* was shifted primarily to the upper small intestine, while still being present in the lower regions of the gut where WT *Pvu* is prevalent. Thus, the context-dependent requirement for DifS may reflect differences in the selective pressures encountered across the length of the gastrointestinal tract. More broadly, the observation that the fitness contribution of the *dif* locus becomes apparent only during direct intraspecies competition highlights the importance of microbial context in determining the fitness consequences of genetic variation. When these *Pvu* strains were introduced sequentially rather than simultaneously, the strain colonizing first consistently maintained a competitive advantage over the strain introduced second, regardless of genotype. These findings are consistent with the concept of priority effects, in which the timing of colonization and interactions with resident bacteria influence bacterial establishment within microbial communities^72,73^.

Although the RNA-seq data indicated that deletions of genes in the *dif* locus led to large-scale changes in cell wall and membrane activities, untargeted lipidomics did not reveal significant differences in overall membrane lipid composition or individual lipid features between the *dif* deletion strains and WT *Pvu*. However, untargeted metabolomics methods identified significant differences in a range of specific metabolites and metabolite classes. In particular, the reduction of a subset of lyso-PE, lyso-PC, lyso-PS, and monoacylglycerol species in Δ*difST* is consistent with potential alterations in turnover or synthesis of membrane phospholipids. These findings suggest that the *dif* deletion strains do not undergo broad lipid remodeling during *in vitro* growth under the defined media conditions tested but may exhibit more localized changes in cellular metabolism. Notably, consistent with the observed transcriptional shifts, the *dif* deletion strains were more sensitive than WT *Pvu* to a variety of antibiotics and bile salts, indicating that measurable changes in bacterial fitness occurred in the absence of *dif* genes, despite a lack of broad alterations to the cellular lipidome. The deletion strains did not exhibit growth defects in BMM but showed reduced growth under BHIS conditions. The absence of antibiotic-associated growth defects in BMM may reflect the substantially slower growth of *Pvu* under these conditions, as many cell wall-targeting antibiotics exert their effects primarily on actively growing cells, and slower growth can reduce susceptibility to these antibiotics^74^. Although the *dif* deletions strains exhibited increased sensitivity to bile salts *in vitro,* Δ*difS* was readily detected in the small intestine using RT-PCR. This was true despite bile acids being present at substantially higher levels in the small intestine compared to the distal gut^75,76^. Thus, bile sensitivity alone is unlikely to account for the altered gastrointestinal distribution of the Δ*difS* deletion strain compared to WT *Pvu*, suggesting that additional environmental factors lead to their compartmentalization to distinct regions of the gut.

Although DifS and its homologs have not previously been characterized in the *Bacteroidaceae* family, a DifS homolog (locus tag: M0N98_01539; hypothetical protein) in another *Pvu* strain, CL10T00C06, was among the top 25 genes associated with fitness in an RB-TnSeq screen performed in a dextran sodium sulfate (DSS) colitis mouse model^22^, although this gene was not discussed by the authors. The independent identification of a DifS homolog in a separate *Pvu* strain and inflammation mouse model nevertheless provides additional support for the role of this protein family in bacterial fitness in the mammalian gut. Together with our findings, these observations establish that DifS-family proteins represent a previously unrecognized component of fitness across *Phocaeicola* within diverse intestinal environments.

Our work identified DifH as a key component of the *dif* locus, and as a member of the HU/IHF superfamily of DNA binding proteins^66^ with a conserved HU-HIG^66^ and winged helix-like domain. Overall, IHF proteins are similar in structure and sequence to HU proteins and belong to the prokaryotic histone-like DNA-bending protein family^77^. Both HU and IHF proteins can affect the transcription of genes by playing both the role of a repressor and activator^77^. Whereas HU proteins are thought to bind DNA in a sequence promiscuous manner, IHF proteins are thought to bind with greater sequence specificity^77^. This relationship reflects their evolutionary history: IHF arose from a lineage of HU proteins, and the residues responsible for its sequence-specific binding were acquired gradually over time^78^. In contrast to the well-characterized Proteobacterial HU/IHF proteins, comparatively little is understood about them in the *Bacteroidetes*. Whereas canonical HU and IHF proteins are known to bind DNA as heterodimers, it remains unknown if DifH binds as a homodimer or heterodimer with an unknown partner, or the extent of functional redundancy with other members of the expanded and rapidly evolving IHF family in the *Bacteroidetes*^66^. Our finding that DifH binds and regulates the expression of one of its close gene homologs, *peg.598*, suggests a degree of regulatory interaction within this family. Notably, *Bacteroidetes* HU/IHF proteins exhibit limited sequence similarity to the more extensively characterized IHF proteins found in *Proteobacteria*, *Firmicutes*, and *Actinobacteria* phyla^77,79^, and form a distinct IHF-like clade, similar to the previously described IHF proteins^78^. Our phylogenetic analysis places DifH within that divergent group of *Bacteroidetes* HU/IHF proteins (Supplemental Figure S6)^66,77^.

We found that DifH binds DNA upstream of its own gene within the *dif* locus. Like the canonical IHF proteins in *E. coli* (IHFα and IHFβ), DifH recognizes a specific 30-35 bp binding motif and binds to hundreds of genomic loci^80,81^. However, these identified HU and IHF motifs are not overtly similar to the DifH motif identified here. Beyond the primary DifH binding motif predicted by bioinformatic analysis, we discovered that the DifH motif occurred in a larger repetitive organization at sites of DifH occupancy: GTGA(10-bp)TCAC(4-bp), 22-bp total, with multiple repeats per peak (Figure 7H and Supplemental Figure S9). While bioinformatic prediction centered the motif on the GTGA(10-bp)TCAC unit, the relaive importance and centrality of the 10-bp and 4-bp gaps remains unknown. The repeating palindromic structure is reminiscent in organization and size to Repetitive Extragenic Palindrome elements (REPs, also called Bacterial Interspersed Mosaic Elements or BIMEs) found in extragenic regions usually in RNA 3’UTRs^82^, to cooperative operator arrays that regulate gene expression in bacterial and phage genomes^83,84^, to iteron arrays controlling plasmid replication^85^, and other examples of complex repetitive DNA elements in bacteria^86,87^. This strict repetitive organization also raises the possibility of high-order multimerization, and the array periodicity (22-bp) is close to the length of two DNA helical turns which may place successive dimers on the same DNA face. The immediate adjacency of these arrayed repeats combined with the predicted DNA bending properties of DifH could cause coiling of that DNA into filaments, which may in turn restrict DNA supercoiling or looping that may affect accessibility and activity of neighboring promoters^88,89^.

Our data indicate that the motif structure is conserved not only in *Pvu* but across other *Bacteroides* strains suggesting a broader role for DifH and its homologs among this bacterial clade. DifH homologs across *Phocaeicola* and *Bacteroides* species are frequently located adjacent to a helicase gene, suggesting a potential functional association between DifH and the neighboring helicase. Prior studies have implicated helicase-associated systems in transcriptional regulation^71^ and in DNA defense against phages^72,73^, and have further shown that IHF proteins regulate bacterial virulence genes and that IHF deletion leads to widespread transcriptional changes and impaired virulence-associated traits. Additionally, the closest DifT homolog in *E. coli*, FadL, is known to be a major mediator of phage sensitivity^90,91^, suggesting potential additional defense functions for the *Pvu dif* locus. These findings support a role for DifH as a DNA-binding transcriptional regulator that modulates specific transcriptional responses rather than being a global regulator.

## ACKNOWLEDGEMENTS

The authors are grateful to Kristen Cleveland, Brittany McAdams, Cassandra Jimenez, Jameson Cameron, Connor Herron, and Molly McCauley in Duke University’s Division of Laboratory Animal Resources (DLAR) for help with gnotobiotic experiments; to Lauren Davey for advice during Tn-seq library preparation; to Todd Bradley for use of the Eppendorf EpMotion liquid handling robot; to Colin Lickwar and Courtney Rivet-Noor for help with ChIP-seq analysis and manuscript editing, respectively.; to Courtney Price for assistance interpreting the LC-MS/MS data.; and to Madeline Colley and Jeffrey Spraggins for guidance on lipidomic analysis.

## DATA AVAILABILITY

All data generated during this study will be deposited in appropriate public data repositories and made publicly available upon publication of this work in a peer-reviewed journal. Accession numbers will be provided at that time. In the interim, data may be requested from the corresponding author.

## DECLARATION OF GENERATIVE AI USE

ChatGPT (OpenAI, GPT-6) was used for copyediting to improve the clarity and grammar of the manuscript. The authors reviewed and edited all AI-assisted text and take full responsibility for the accuracy and integrity of the final manuscript.

