## Supplemental Figure Legends for "A microbiota-dependent fitness locus promotes intraspecies competition and cell envelope resilience in gut bacterium *Phocaeicola vulgatus*"

### SUPPLEMENTARY FIGURE LEGENDS

**Supplementary Figure S1: Construction, validation, and analysis of the *Pvu* transposon library.**
**(A)** Southern blot analysis of randomly selected transposon mutants confirming predominantly single transposon insertions within the arrayed library.
**(B)** Correlation of transposon insertion counts per gene between two independently sequenced input library aliquots.
**(C)** Genome-wide distribution of transposon insertions across the *Pvu* CL09T03C04 genome.
**(D)** Rarefaction analysis showing sequencing depth and unique transposon insertion sites for the input library and all *in vivo* samples.
**(E)** Heatmap of log2FC values relative to input library for genes represented in the transposon library across all samples. Samples are clustered by time point, with GF and SPF mice fecal samples pooled within each time point. *Pvu* genes represent pooled transposon mutants mapping to the same gene.
**(F)** Principal component analysis (PCA) of untargeted LC-MS/MS metabolomic profiles from selected transposon mutants and WT *Pvu* grown in BHIS (n = 5), based on Euclidean distances. Pairwise PERMANOVA on the same distance matrix indicated significant separation from WT for *Pvu255::Tn* (F = 9.19, $R^2$ = 0.535, BH-adjusted p = 0.0357) and *Pvu2207::Tn* (F = 18.1, $R^2$ = 0.694, BH-adjusted p = 0.0200). *Pvu1650::Tn* and *Pvu32::Tn* did not differ significantly from WT after Benjamini-Hochberg correction for multiple comparisons.
**(G)** Heatmap of the top 30 differentially abundant metabolites in selected transposon mutants relative to WT *Pvu* grown in BHIS, identified using an FDR Benjamin–Hochberg correction (adjusted p-value<0.2) threshold from pairwise comparisons and clustered by metabolite abundance patterns.
**(H)** Transposon insertion map of the *Pvu2206-Pvu2208* locus showing TA insertion sites and mutant abundance in the input library and throughout the GF and SPF mouse experiments.

**Supplementary Figure S2: Genetic and structural characterization of the *Pvu2206–Pvu2208* locus**
**(A)** RT-PCR validation of the *Pvu2206-Pvu2208* locus in WT *Pvu* and mutant strains for confirmation of gene deletions.
**(B)** RNA-seq track from WT *Pvu,* grown in BHIS and BMM, showing the *Pvu2206-2208* locus. The top track shows a zoomed-in view of the locus, while the bottom track displays the full transcript height.
**(C-D)** Growth curves of WT *Pvu* and the *PvuΔ2206, PvuΔ2207, PvuΔ2208*, and *PvuΔ2207-2208* strains (n = 12) in BHIS **(C)** and BMM **(D)**.
**(E)** PONDR disorder protein score for Pvu2207 using the VL-XT algorithm in which the solid line represents the disorder score for each amino acid residue. Any residues above the 0.5 horizontal threshold are predicted to be structurally disordered.
**(F)** AlphaFold structural predictions and confidence scores for Pvu2206, Pvu2207, and Pvu2208.
**(G)** AlphaFold predicted aligned error (PAE) plots for the five ranked models showing a well-defined Pvu2207 N-terminal core.
**(H)** Structural prediction of the core of Pvu2207 consisting of residues G63–Y187, identified using PONDR, IUPred3, and Alphafold shown as the dashed box in **(F)**, visualized in UCSF ChimeraX.
**(I-J)** Growth curve of WT *Pvu* and *PvuCL09difS-FLAG* (n = 8) in BHIS **(I)** and BMM **(J).**
**(K)** Western blot of WT *Pvu* and *PvuCL09difS-FLAG* cells and supernatant fractions probed with an anti-FLAG antibody. The FLAG-tagged DifS protein was detected at the expected molecular weight of approximately 50 kDa (P= pellet, S= supernatant).

**Supplementary Figure S3: Genome-wide transcriptional responses to *difS* and *difH* deletion across in *vitro* growth conditions.**

**(A)** PCoA of RNA-seq samples from WT *Pvu,* Δ*difS*, and Δ*difH* separated by growth condition (BHIS and BMM).
**(B)** Normalized transcript counts of genes within the *dif* operon in WT *Pvu* grown in BHIS and BMM.
**(C)** Heatmap showing genome-wide log2 fold-change values for Δ*difS* and Δ*difH* with pooled replicated (n = 3) compared with WT *Pvu* across BHIS and BMM conditions.
**(D)** Venn diagram showing the number of significantly upregulated and downregulated genes identified for each deletion strain and growth condition.
**(E-F)** Scatterplots showing genome-wide differential expression of Δ*difS* **(D)** and Δ*difH* (**E**) compared with WT *Pvu* in BHIS and BMM. Each point represents a gene, with significant differentially expressed genes highlighted and genes within the *dif* operon indicated in red.
**(G-I)** COG enrichment analysis showing significantly altered functional categories when comparing BMM-grown samples to BHIS-grown samples for WT *Pvu* **(G),** Δ*difH* **(H),** and Δ*difS* **(I).**
**(J-K)** COG enrichment analysis showing significantly altered functional categories in Δ*difH* **(J)** and Δ*difS* **(K)** compared with WT *Pvu* under BMM and BHIS conditions.
Category enrichment was determined using a one-sided Fisher's exact test. Significance: adjusted p values *, < 0.05; **, < 0.01; ***, < 0.001.

**Supplementary Figure S4: Lipidomics and metabolomics profiling reveals changes in select metabolites**
**(A-B)** PCA plot of all clean deletion strains that underwent untargeted lipidomics **(A)** and untargeted metabolomics **(B)** based on Euclidean distances.
**(C-E)** The membrane composition of WT and *dif* deletion strains as depicted through the major membrane lipid classes **(C)**, sphingolipid class **(D)**, and glycerophospholipid class **(E).**
**(F)** Heatmap of all significant metabolites detected through untargeted metabolomics using log2FC when compared to WT using FDR Benjamin–Hochberg correction (adjusted p-value < 0.2).

**Supplementary Figure S5: Growth responses of WT *Pvu* and *dif* deletion strains to membrane and intestinal stressors**
Growth curves of WT *Pvu* and the indicated *dif* deletion strains (n = 3) in the presence of four concentrations of polymyxin B **(A),** carbenicillin **(B),** meropenem **(C),** fosfomycin **(D),** sodium deoxycholate **(E),** TCA **(F),** cholic acid **(G),** and 1:1 sodium deoxycholate:cholic acid **(H).**

**Supplementary Figure S6: Phylogenetic analysis of DifH and IHF/HU-HIG-like proteins**
**(A)** Phylogenetic tree of DifH, homologous proteins, and distantly related proteins from 17 *Bacteroidaceae* type strains identified by BLASTP (E-value ≤ 1), alongside representative HU/IHF superfamily proteins.
**(B)** Zoomed in region of the phylogenetic tree to observe DifH and its closely related proteins.

**Supplementary Figure S7: Validation of the *PvuCL09difH-HA* strain used for ChIP-seq**
**(A)** Western blot of WT *Pvu* and the *PvuCL09difH-HA* strain probed with an anti-HA antibody. The HA-tagged DifH protein was detected at the expected molecular weight of approximately 25 kDa.
**(B)** RT-PCR analysis of the *dif* operon and neighboring genes in WT *Pvu* and the *PvuCL09difH-HA* strain. The red box indicates the HA tag fused to the C-terminus of *PvuCL09difH-HA.*
**(C-D)** Growth curve of WT *Pvu* and *PvuCL09difH-HA* (n = 3) in BHIS **(C)** and BMM **(D).**
**(E)** Competitive fitness of the *PvuCL09difH-HA* strain following co-administration with WT *Pvu* into SPF mice. Relative abundance is shown as the *PvuCL09difH-HA*/WT ratio.

**Supplementary Figure S8: Validation of *PvuCL09difH-HA* ChIP-seq peaks and identification of DNA-binding motifs**

**(A-B)** Summary plot and heatmap of pooled WT *Pvu* and *PvuCL09difH-HA* ChIP-seq replicates grown in BHIS **(A)** and BMM **(B).** Representative peaks identified by MACS2 from the top, middle, and lower portions of the heatmap are shown to illustrate peak quality based on log2 fold enrichment of *PvuCL09difH-HA* relative to WT *Pvu* and input.
**(C-D)** Peak summit enrichment for all ChIP-seq peaks identified by MACS2 ($q < 0.001$) from *PvuCL09difH-HA* and input samples grown in BHIS **(C)** and BMM **(D).** Peaks containing the MEME-identified DNA-binding motif are highlighted in blue.
**(E-F)** Distribution of MEME-identified DNA-binding motifs relative to ChIP-seq peak summits in BHIS **(E)** and BMM **(F).** Motif positions are shown within ±500 bp of each peak summit.
**(G)** Frequency of the BHIS DNA-binding motif in the genomes of *Pvu* and closely related *Phocaeicola* and *Bacteroides* species.

**Supplementary Figure S9: Analysis of DifH-binding motif architecture.**
DNA sequences surrounding the identified DNA-binding motif at each genomic location where DifH binding was detected in both BHIS and BMM. The motif consists of two highly conserved sequences, GTGA (gray boxes) and TCAC (black boxes), separated by 10 bp, as indicated by solid lines. Successive motif units are spaced 4 bp apart, as indicated by dashed lines. Boxes with white lines indicate instances in which either conserved sequence contains a mismatch or inversion.
