## Supplementary figures and images for "A microbiota-dependent fitness locus promotes intraspecies competition and cell envelope resilience in gut bacterium *Phocaeicola vulgatus*"

### Supplementary Figure S1

#### Supplementary Figure S1

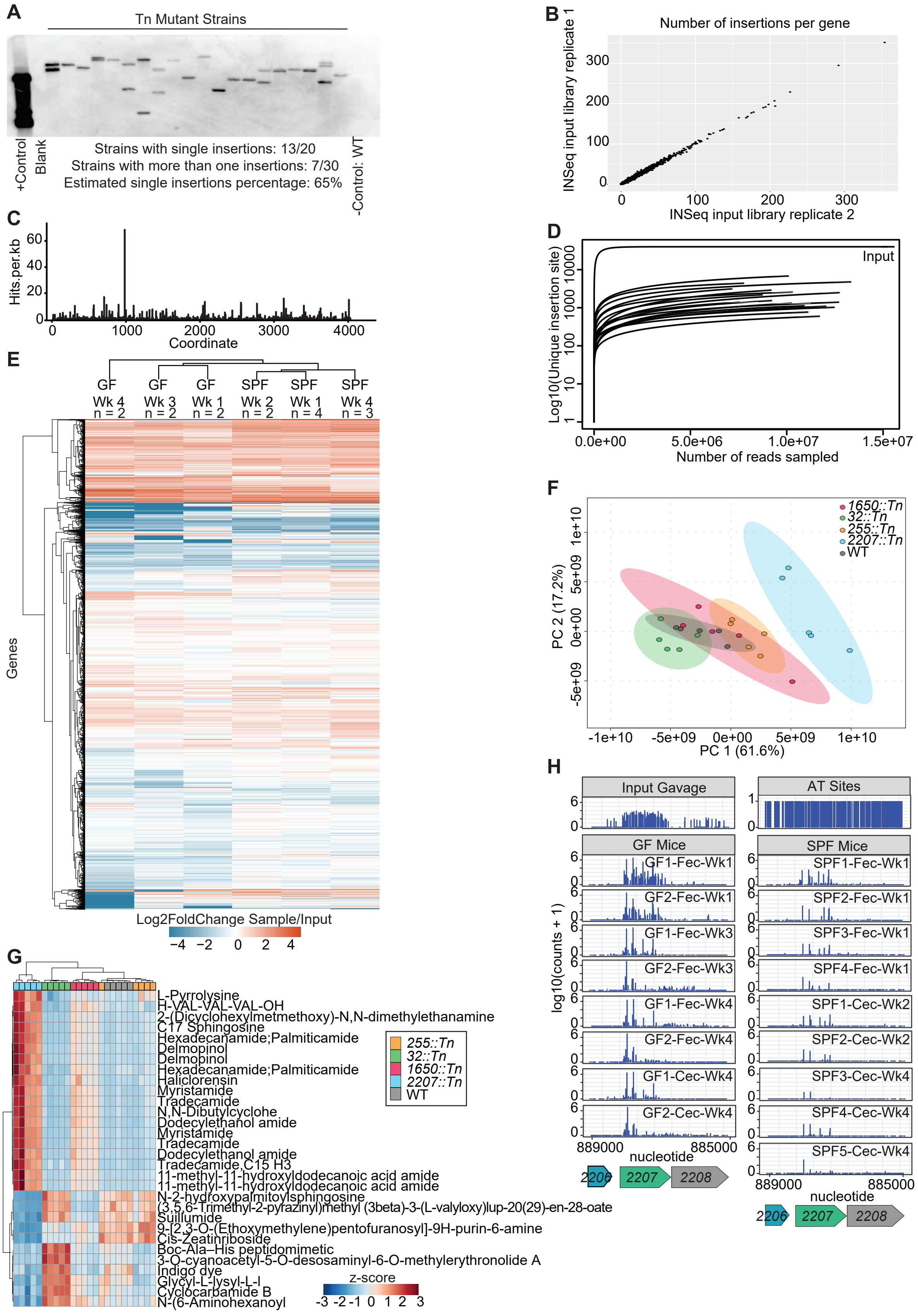

### Supplementary Figure S2

#### Supplementary Figure S2

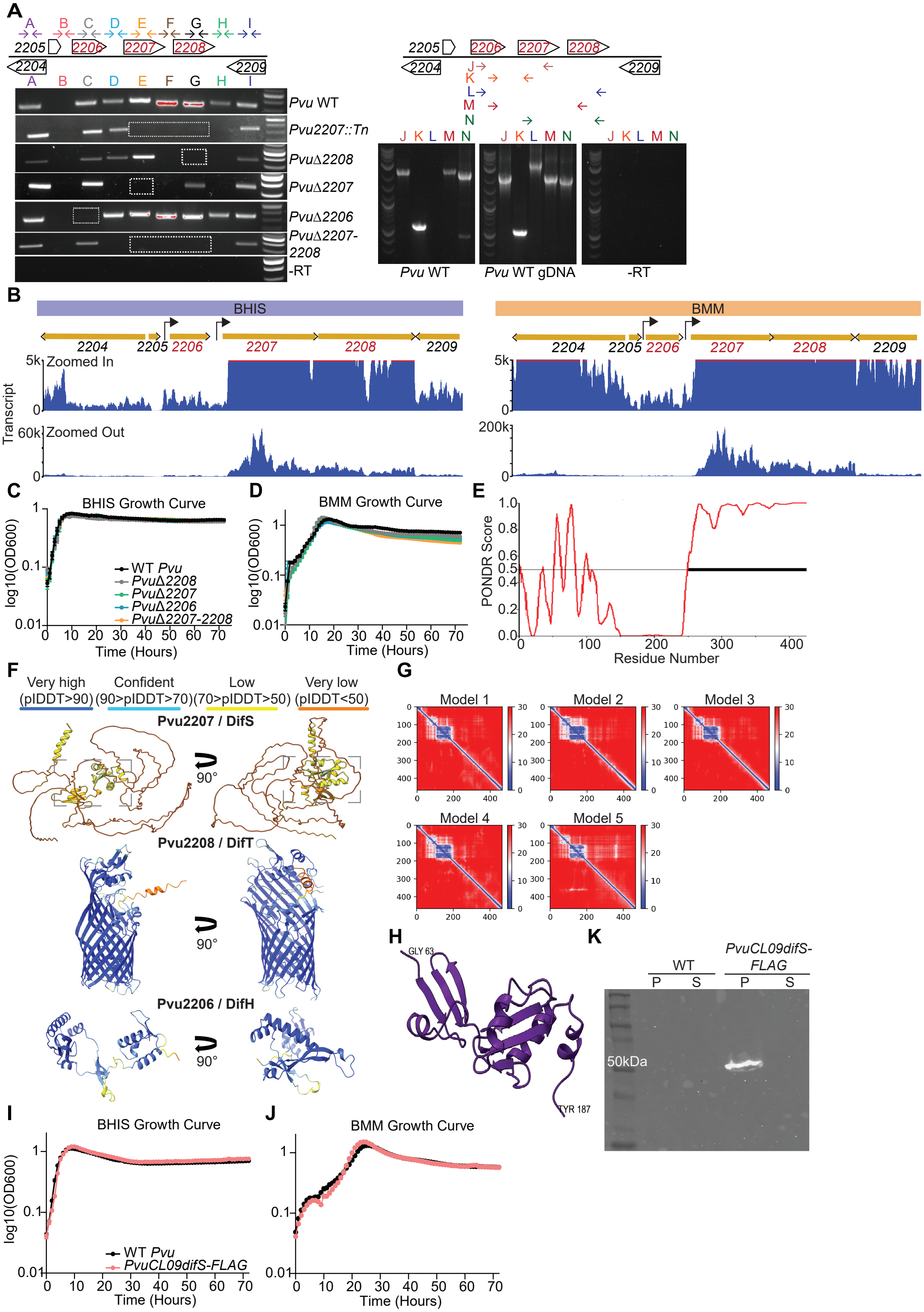

### Supplementary Figure S3

#### Supplementary Figure S3

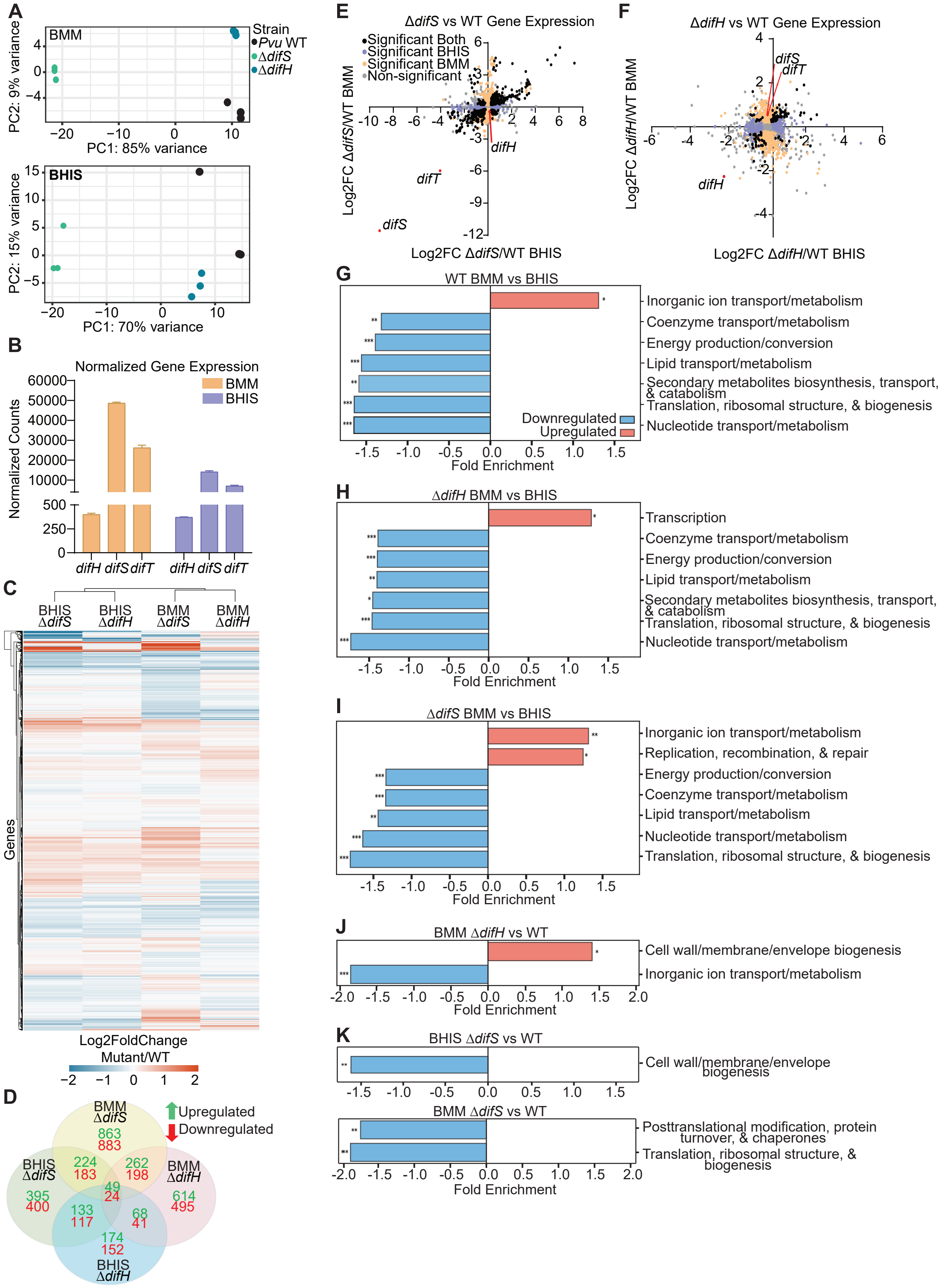

### Supplementary Figure S4

#### Supplementary Figure S4

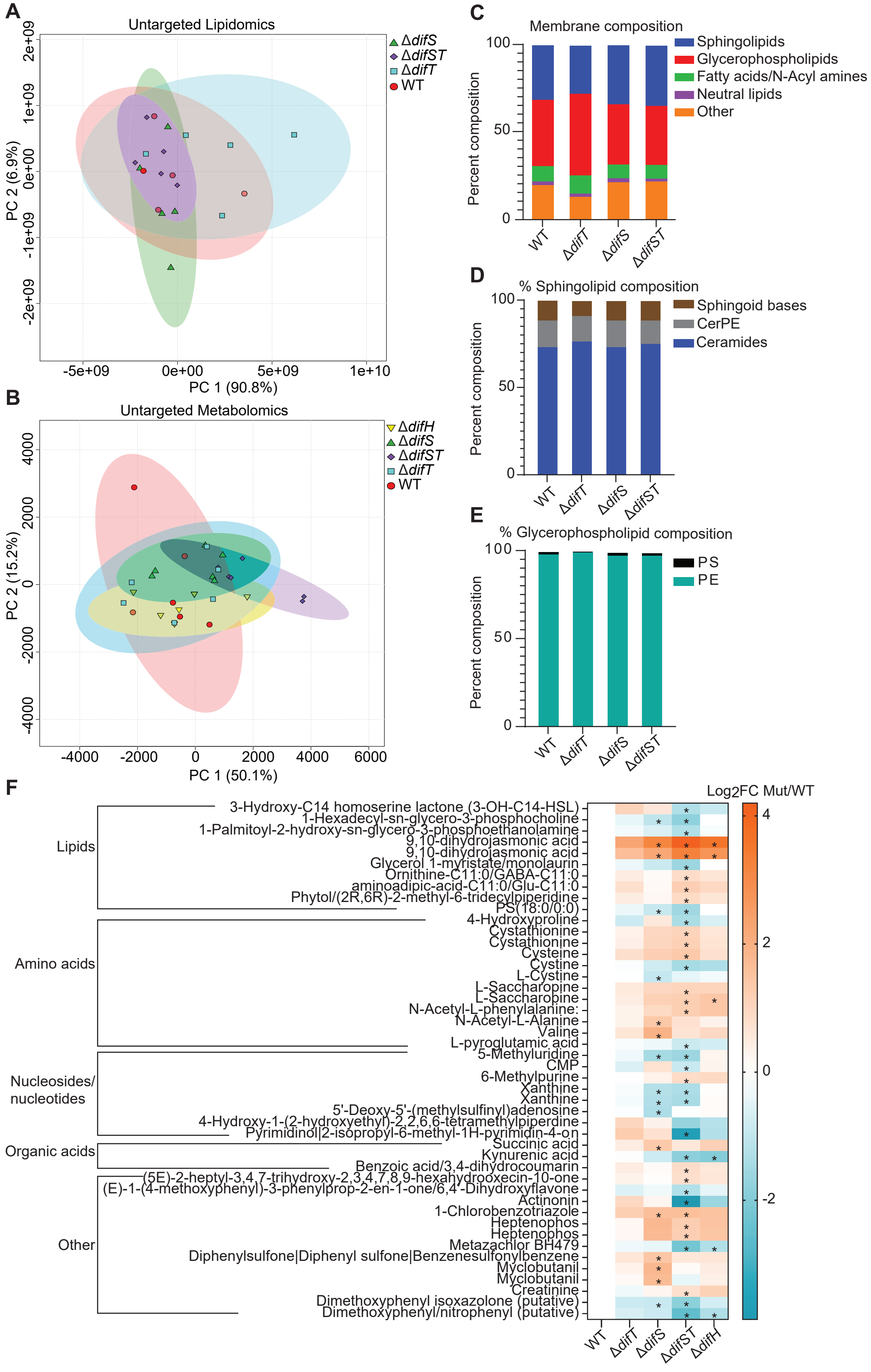

### Supplementary Figure S6

#### Supplementary Figure S6

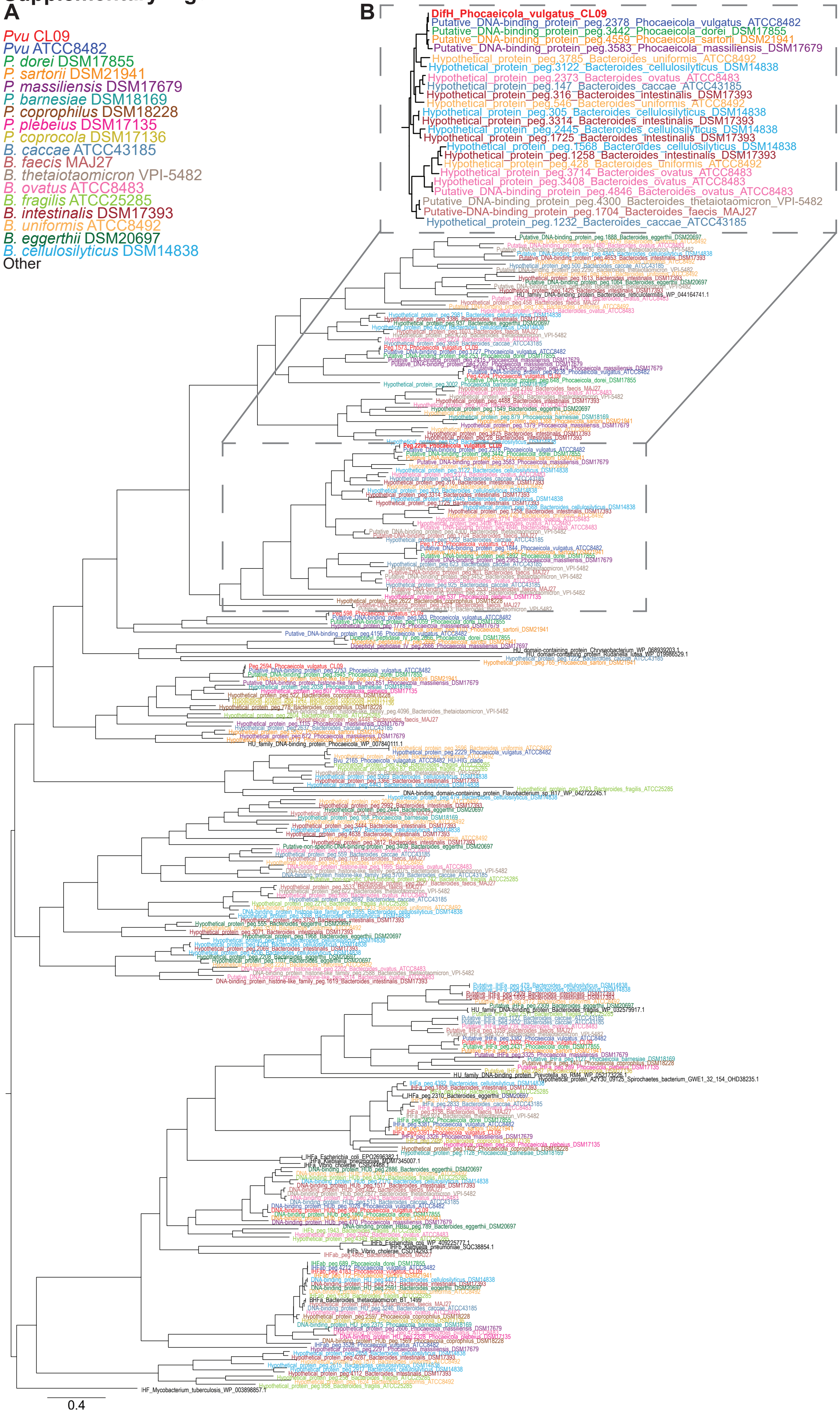

### Supplementary Figure S7

#### Supplementary Figure S7

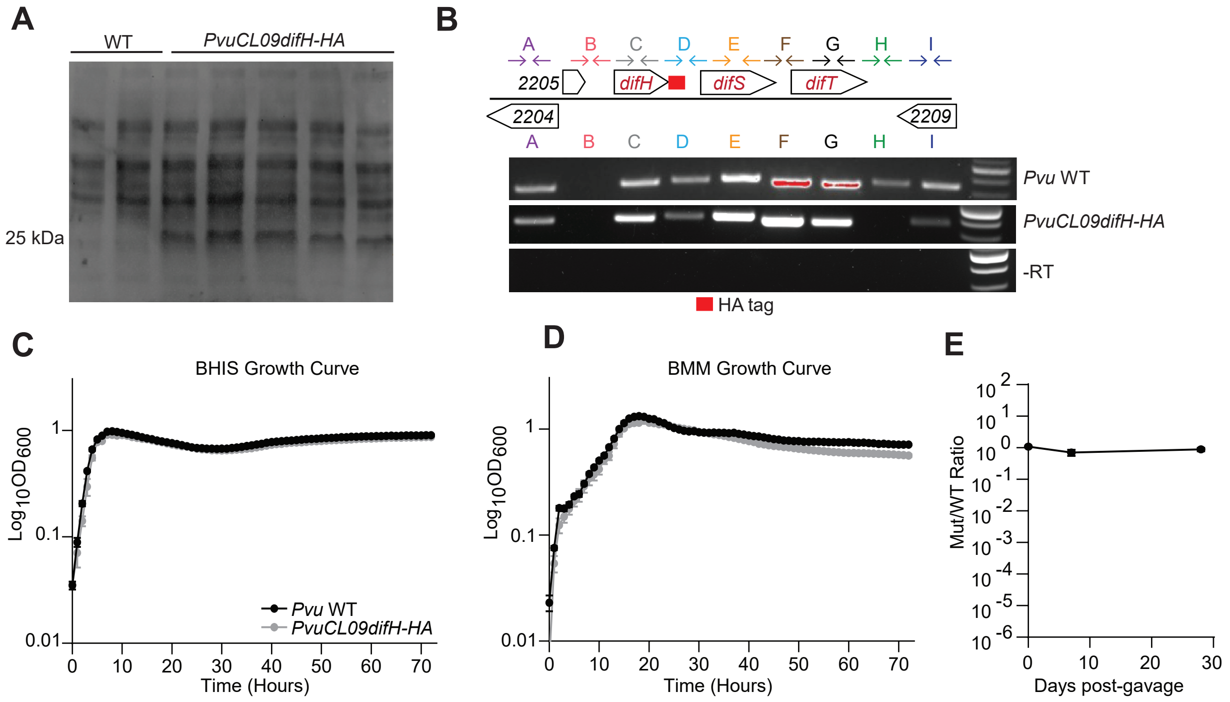

### Supplementary Figure S9

#### Supplementary Figure S9

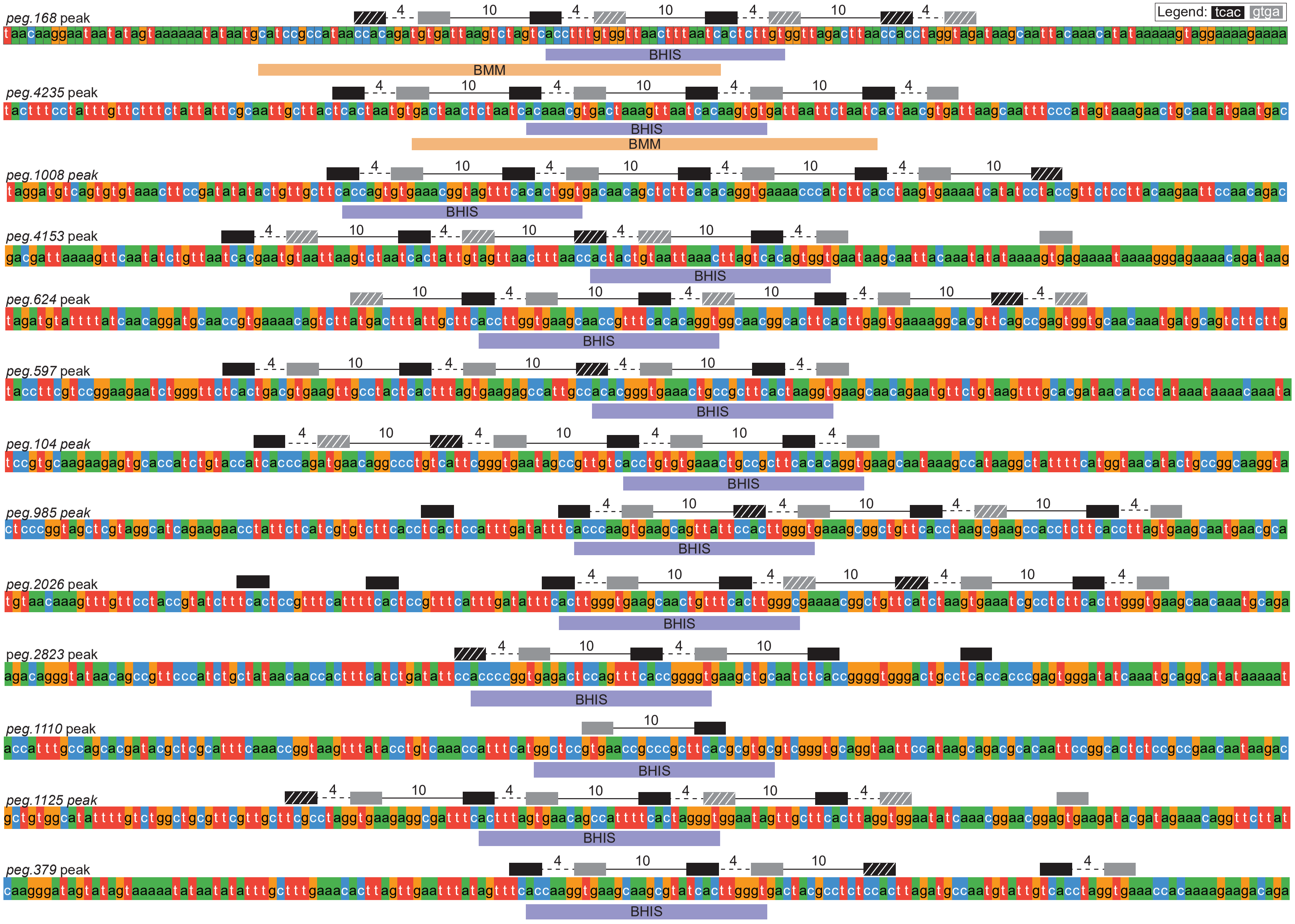
