## Supplementary Figure S5 for "A microbiota-dependent fitness locus promotes intraspecies competition and cell envelope resilience in gut bacterium *Phocaeicola vulgatus*"

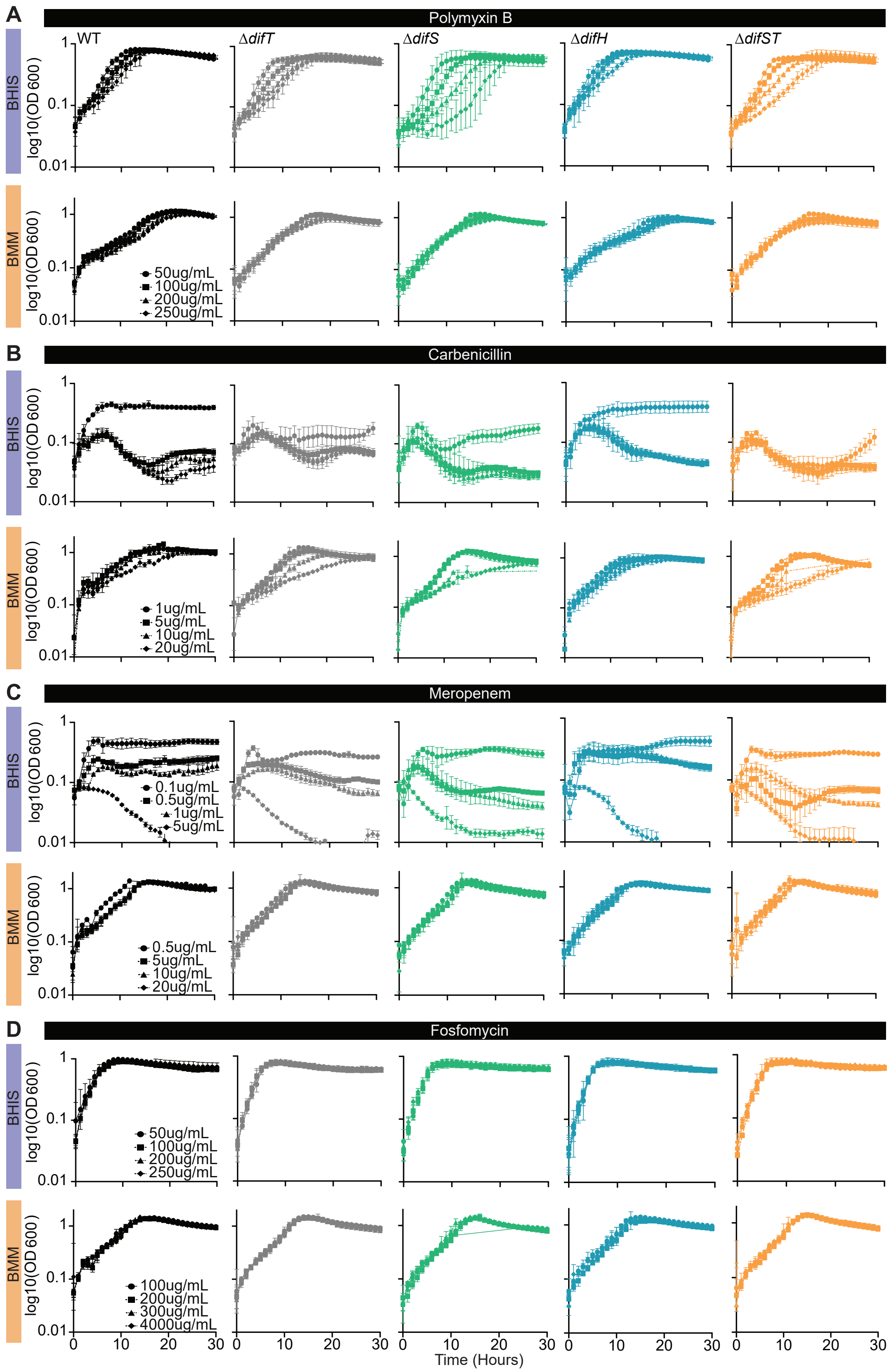

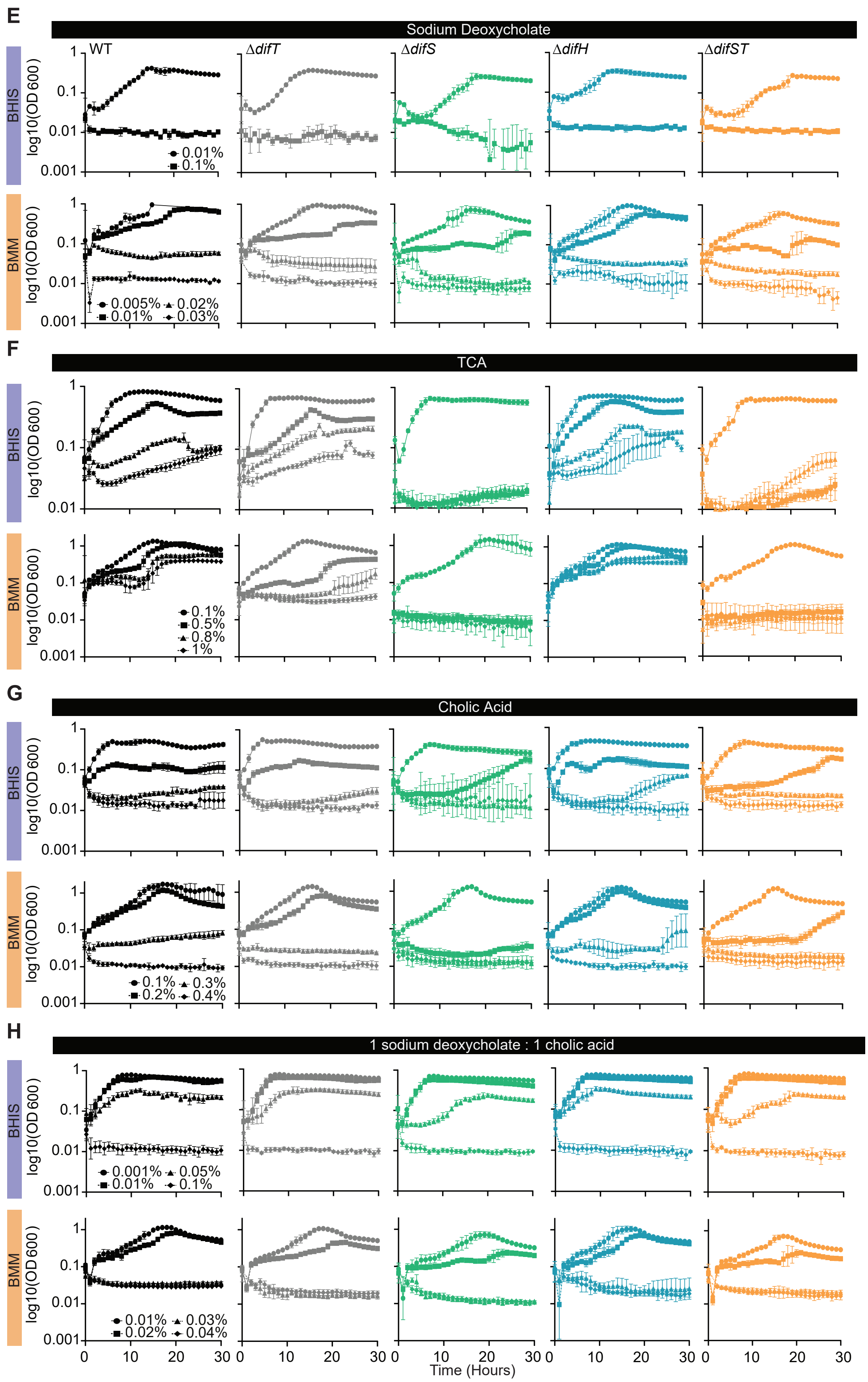
E
Sodium Deoxycholate
WT
ΔdifT
ΔdifS
ΔdifH
ΔdifST
BHIS
BMM
log10(OD 600)
1
0.1
0.01
0.001
0.01%
0.1%
0.005%
0.02%
0.01%
0.03%
F
TCA
0.1%
0.5%
0.8%
1%
G
Cholic Acid
0.1%
0.3%
0.2%
0.4%
H
1 sodium deoxycholate : 1 cholic acid
0.001%
0.05%
0.01%
0.1%
0.01%
0.03%
0.02%
0.04%
0
10
20
30
Time (Hours)
