## Supplementary Figure S8 for "A microbiota-dependent fitness locus promotes intraspecies competition and cell envelope resilience in gut bacterium *Phocaeicola vulgatus*"

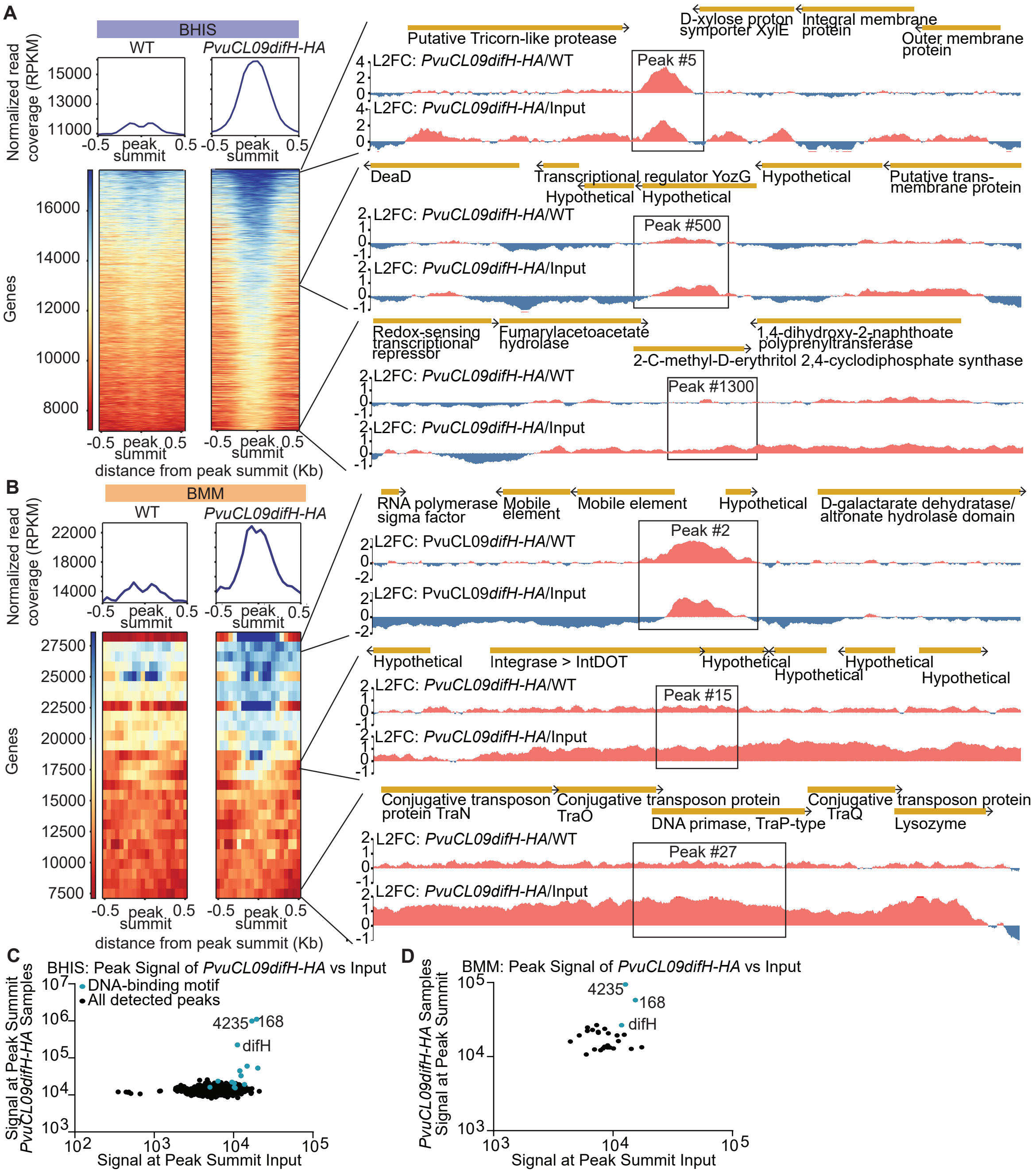


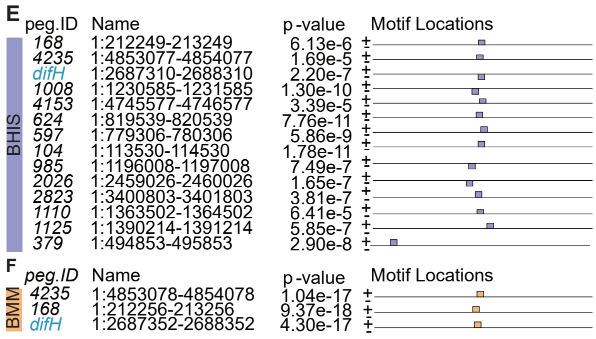


G

| Strain | Unique Motif Hits | Genome size (Mb) | Motifs per Mb |
|---|---|---|---|
| *Pvu* CL09 | 383 | 4.88 | 78.5 |
| *Pvu* ATCC 8482 | 372 | 5.16 | 72.0 |
| *B.barnesiae* | 294 | 3.62 | 81.3 |
| *B.caccae* | 323 | 4.57 | 70.7 |
| *B.cellulosilyticus* | 450 | 6.87 | 65.5 |
| *B.coprocola* | 264 | 4.30 | 61.4 |
| *B.coprophilus* | 233 | 4.04 | 57.7 |
| *B.eggerthii* | 233 | 4.20 | 55.5 |
| *B.faecis* | 414 | 6.11 | 67.7 |
| *B.fragilis* | 379 | 5.24 | 72.3 |
| *B.intestinalis* | 462 | 6.05 | 76.4 |
| *B.ovatus* | 446 | 6.47 | 68.9 |
| *B.theta* | 427 | 6.29 | 67.8 |
| *B.uniformis* | 319 | 4.63 | 68.9 |
| *P. massiliensis* | 313 | 4.51 | 69.4 |
| *P.dorei* | 416 | 5.57 | 74.7 |
| *P.plebeius* | 324 | 4.42 | 73.3 |
| *P.sartorii* | 388 | 5.29 | 73.3 |
