## Supplemental Methods for "A microbiota-dependent fitness locus promotes intraspecies competition and cell envelope resilience in gut bacterium *Phocaeicola vulgatus*"

### *Pvu* genome sequencing

Pre-existing assembled short sequencing reads obtained via Illumina sequencing were downloaded from the European Nucleotide Archive (SRR357374)[1]. Long read sequencing was performed at SeqCenter using Oxford Nanopore sequencing on *Pvu* CL09T03C04 gDNA. Quality control and adapter trimming were performed using porechop[2] (v0.2.3) and the assembly was downloaded as a BAM file. Long sequencing reads were assembled using the Flye program[3] (v2.8.2) using default settings and the "--plasmids" attribute to yield a single, non-circular contig with no plasmids. This assembly was supplied to the Unicycler program (v0.4.8) and combined with the short-read sequencing data described above to yield a circular, resolved genome for *Pvu* CL09T03C04 using the attributes "--exisiting_long_read_assembly" and "--mode conservative". The assembled genome was annotated using the RASTtk tool kit as implemented on the BV-BRC bioinformatics database[4,5]. The resolved version of the genome is available at NCBI [submission in progress].

### Sequencing arrayed *Pvu* Tn mutant library clones

Aliquots of the frozen *Pvu* transposon library were swabbed onto pre-reduced tryptic soy agar (TSA) plates supplemented with 5% sheep's blood (VWR, #10324-332) and incubated anaerobically at 37˚C for 48 hours. Colonies were picked from the TSA plates directly into forty 1mL deep well 96-well plates (USA Scientific, #1896-1800) containing pre-reduced TYG medium supplemented 5µg/mL erythromycin, 50µg/mL gentamicin, and 0.5g/L cysteine, leaving the last well on each plate empty to serve as a blank for a total of approximately 3,000 arrayed *Pvu* transposon mutants. The deep well plates were incubated anaerobically at 37˚C overnight, after which 200µL of culture was taken from each well and transferred to a sterile 96-well culture plate to serve as a replicate to be used for combinatorial pooling. All plates were frozen at -80˚C after the addition of sterile 50% glycerol solution.

Combinatorial pooling was performed on thawed 96-well plates using an EpMotion 5075 Automated Pipetting System (Eppendorf), according to previously described protocols[6], to create 24 combinatorial pools to be used for high-throughput sequencing.

*Pvu* genomic DNA was extracted from the 24 combinatorial pools using the DNeasy Blood and Tissue Kit (Qiagen, # 69504) and eluted into 200µL of elution buffer. Genomic DNA from each pool was used for high-throughput sequencing library preparation using previously described protocols[6], with modified sequencing barcodes (Supplemental Table S2) for compatibility with the updated Illumina HiSeq 4000 sequencer. High-throughput sequencing was performed at the Duke Center for Genomic and Computational Biology. Arrayed library and mouse Tn-seq samples were sequenced on a HiSeq4000 and NovaSeq 6000 single lane, respectively, with a 50bp amplicon and 15% PhiX spiked in.

### RT-PCR of the *dif* locus

Cells were grown overnight in BHIS anaerobically at 37˚C, subcultured to an O.D.600 of 0.5-0.6 followed by treatment with 2:1 v:v of RNAprotect Bacteria Reagent (Qiagen, #76506) for 5 minutes at room temperature. RNA was extracted from subsequent pellets using the Roche High Pure RNA Isolation kit (#11828665001). RNA was subjected to DNAseI treatment following Invitrogen DNaseI treatment followed by cDNA synthesis (ProtoScript® II First Strand cDNA Synthesis Kit, #E6560). RT-PCR was done using the cDNA using GoTaq Master Mix (Promega, Cat# M7123) and a general 3 step PCR protocol was adjusted for 45 cycles and a 6 minute elongation step. See Supplementary Table S8 for all the primer pairs that were used for RT-PCR.

**Generation and validation of clean deletion mutants**

All *Pvu* deletion strains were generated by conjugation-mediated plasmid transfection and selection of double-crossover resolvents with an anhydrotetracycline (aTC)-inducible ssBfe1 cassette contained in the vector pLGB13[7] (Addgene, plasmid #126618). Sequences of approximately 1,000bp in length flanking the appropriate coding regions were amplified by PCR and assembled into the EcoRV site of the pLGB13 vector using Gibson Assembly (NEB, Cat# E2611S). 3µL aliquots of each assembly reaction was transformed into competent donor *Escherichia coli* PIR2 (Thermofisher, # C111110). Assembled plasmids were validated via sequencing (Plasmidsaurus) and then transformed into *Escherichia coli* S17 λ*pir* (a gift from Raphael Valdivia) and conjugated into *Pvu* as described previously[7]. The first crossover event was confirmed by PCR with gene specific primers (Supplementary Table S8). Double cross over events were selected on 40ng/mL of aTC and incubated anaerobically at 37˚C for 36 hours. Mutant clones were identified by PCR screening.

RNA extraction was performed as described above. Growth curves of all mutants were performed in a Spectrostar Nano (BMG Labtech) placed in a Coy anaerobic chamber in BHIS and BMM alongside WT *Pvu.* Data was then analyzed and graphed using GraphPad Prism version 11.0.2.

**Conservation analysis of *Pvu* and its close congeners**

The whole genome sequence phylogenetic tree was generated using Bacterial Genome Tree tool in BV-BRC[4] and the tree was visualized using FigTree[8] v1.4.4. The gene neighborhoods and homologs of the *dif* genes were observed using Fast.genomics[9]. Homologs of DifS and Pvu2206 were identified using BLASTP in BV-BRC (e-value < 1). DifH homologs were identified using BLASTP searches against the evaluated Bacteroidaceae genomes. Because numerous DifH-related proteins were identified, hits were categorized based on amino acid sequence similarity to the Pvu DifH protein. Proteins sharing >40% amino acid similarity with DifH were classified as “close homologs”, whereas proteins sharing 20–40% amino acid similarity were classified as “distant homologs”. Proteins with <20% amino acid similarity were not included in the homolog classification. The phylogenetic tree of DifS and its homologs was generated using BV-BRC; however, since the phylogenetic tree of DifH, its homologs, canonical IHF proteins (sequences obtained from GenBank), and other previously described proteins[10] had too large of a FASTA file, the sequences were first aligned using CLUSTAL Omega[11] and then generated into a tree in BV-BRC. All trees were visualized using FigTree v1.4.4.

***In vitro* RNA-seq and analysis**

WT *Pvu* (n =3), Δ*difH* (n =3), and Δ*difS* (n =3) cultures grown overnight in BHIS and BMM anaerobically at 37°C, subcultured 1:100 and grown to O.D.600 of 0.5-0.6. RNAprotect Bacteria Reagent (Qiagen, #76506) was then added at a 2:1 v:v ratio and incubated at room temperature for 5 minutes before following the RNA extraction methods described above. RNA samples were quantified by Qbit (Thermofisher, # Q32855), treated with DNAseI (Sigma, #D4513-1VL) and rRNA depletion was performed using custom *Pvu*-specific Ribozero probes (Illumina). Samples were DNAse treated with Invitrogen DNAse (RNAse free). Library preparation was performed using Illumina’s Stranded Total RNA Prep Ligation with Ribo-Zero Plus kit and 10bp IDT for Illumina indices. Three technical replicates of each strain in each media condition were then submitted for high throughput RNA sequencing (SeqCenter Pittsburgh, PA) for Illumina sequencing on a NextSeq2000 giving 2x51bp reads to generate 12M paired end reads with the *Pvu* CL09T03C04 genome as the reference.

Demultiplexing, quality control, and adapter trimming were performed with bcl-convert (v3.9.3)[12]. Quality control and adapter trimming were performed with bcl2fastq[13]. Read mapping was performed with HISAT2[14]. Read quantification was performed using Subread’s featureCounts[15] functionality. Read counts loaded into R[16] and were normalized using edgeR's[17] Trimmed Mean of M values (TMM) algorithm. Subsequent values were then converted to counts per million (cpm). Differential expression analysis was performed using edgeR's exact test for differences between two groups of negative-binomial counts with an estimated dispersion value of 0.1. Principal component analysis (PCA) plots were generated using the normalized expression values of all genes. Heatmaps were generated using the pheatmap[18] (v1.0.13) and were based on DESeq2[19]-derived (v1.50.2) log2 fold changes and normalized count data. Data visualization was generated using GraphPad Prism version 11.0.2.

Differentially expressed genes (DEGs) were mapped to COG functional categories using eggNOG-mapper[20] to identify significantly enriched COGs in all strains grown in BMM versus BHIS as well as all deletion strains versus WT *Pvu* and were then visualized using ggplot2[21]. Briefly, a functional annotation for the *Pvu* CL09T03C04 closed genome was performed with eggNOG-mapper v3.0.0[22] using orthology assignments from the eggNOG 7 database[23]. Homology searches used DIAMOND in sensitive mode (e-value ≤ 0.001), transferring GO, KEGG, COG and PFAM terms from orthologs within the automatically adjusted taxonomic scope (--tax_scope auto). Significantly enriched or depleted genes without predicted functional annotations were labeled as NA or function unknown. Category enrichment was determined using a one-sided Fisher's exact test to determine if any category was over-represented in the test set relative to the background representation of the relevant category in the dataset. Any category represented by fewer than 2 genes in the test set were excluded from testing to reduce noise from low-count categories. Resulting raw p-values were corrected for multiple comparisons[24].

**LC-MS bacterial culture conditions**

*Pvu* cultures were initiated as described above. Cells were collected by centrifugation at 10,000 × g for 2 min, after which the supernatant was aspirated. Cell pellets were immediately frozen and stored at −80°C until metabolite or lipid extraction.

**Untargeted metabolomics of *in vitro* bacterial cultures**

*Sample preparation:* Frozen bacterial cell pellets were resuspended in 1 mL of 80% LC-MS-grade methanol and vortexed for 5 min. Samples were centrifuged at 10,000 × g for 10 min at 4°C, and the resulting supernatants were filtered through 0.22-µm PVDF membrane filters (VWR, #76018-882). Filtrates were dried using a SpeedVac at room temperature and stored at −20°C until LC-MS/MS analysis.

*Mass spectrometry analysis:* Liquid chromatography-mass spectrometry (LC-MS/MS) data were acquired using a Q Exactive HF-X quadrupole-Orbitrap mass spectrometer equipped with a heated electrospray ionization (HESI) source and coupled to a Vanquish UHPLC system (Thermo Fisher Scientific). Samples (n = 6 per group) were diluted two-fold in 50% acetonitrile prior to analysis. Samples were randomized prior to injection, and 2 µL of each sample was injected onto an Acquity BEH C18 column (1.7 µm, 2.1 × 100 mm; Waters) operated at a flow rate of 0.3 mL/min. Mobile phase A consisted of water containing 0.1% formic acid, and mobile phase B consisted of acetonitrile containing 0.1% formic acid. The gradient began at 99.9% A and 0.1% B and was maintained for 1 min, followed by a linear decrease in A to 85% from 1–5 min, 25% from 5–17 min, and 0.1% from 17–21 min. The mobile phase was maintained at 0.1% A from 21–24 min, followed by a linear return to the initial conditions from 24–24.2 min, which were maintained until 27 min. The first minute of each run was diverted to waste.

Data were acquired in positive ionization mode over a full-scan range of m/z 80–1,200. The capillary temperature was 320°C, tube lens offset was 55 V, and spray voltage was 3.70 kV. Sheath, auxiliary, and sweep gas flows were 35, 10, and 1 arbitrary units, respectively. Full-scan spectra were acquired at a resolution of 60,000 with an automatic gain control (AGC) target of 1E6. Data-dependent MS/MS spectra were acquired for the 10 most intense precursor ions using stepped normalized collision energies of 15, 35, and 60. Precursor ions were dynamically excluded for 5 s following fragmentation. MS/MS spectra were acquired at a resolution of 15,000 with an AGC target of 2E5.

**Untargeted lipidomics of *in vitro* bacterial cultures**

*Sample preparation:* Frozen cell pellets prepared as described above were subjected to organic solvent extraction. Pellets were resuspended in 200 µL of LC-MS-grade water and spiked with 5 µL of cardiolipin-d5 internal standard (Avanti Polar Lipids, #330736; 100 µg/µL). Samples were sonicated on ice with metal beads (Next Advance, #NAVYR1) in a water bath sonicator for 30 min. Following sonication, 200 µL of LC-MS-grade methanol and 800 µL of methyl tert-butyl ether (MTBE) were added. Samples were vortexed for 1 min and centrifuged at 500 × g for 10 min. A 700-µL aliquot of the upper organic phase was transferred to a new vial and dried using a SpeedVac. Dried extracts were stored at −20°C until LC-MS/MS analysis.

*Mass spectrometry analysis:* Lipidomic analyses were performed using the Q Exactive HF-X/Vanquish UHPLC platform described above. Samples (n = 5 per group) were resuspended in 300 µL of 100% methanol prior to analysis. Samples were randomized prior to injection, and 5 µL of each sample was injected onto an Acquity UPLC CSH C18 column (1.7 µm, 2.1 × 50 mm; Waters) operated at a flow rate of 0.25 mL/min. Mobile phase A consisted of 60:40 acetonitrile containing 0.1% formic acid and 10 mM ammonium formate, and mobile phase B consisted of 90:10 isopropanol containing 0.1% formic acid and 10 mM ammonium formate. The gradient began at 100% A and was decreased linearly to 68% A from 0–2 min, 55% A from 2–4 min, 48% A from 4–5 min, 42% A from 5–8 min, 34% A from 8–11 min, 30% A from 11–14 min, 25% A from 14–18 min, and 1% A from 18–19 min. The mobile phase was maintained at 1% A from 19–22 min, followed by a linear return to 100% A from 22–25 min. The first minute and final 7 min of each run were diverted to waste.

Data were acquired in positive ionization mode over a full-scan range of m/z 100–1,350. The capillary temperature was 300°C, tube lens offset was 45 V, and spray voltage was 3.80 kV. Sheath, auxiliary, and sweep gas flows were 50, 10, and 1 arbitrary units, respectively. Full-scan spectra were acquired at a resolution of 60,000 with an AGC target of 1E6. Data-dependent MS/MS spectra were acquired for the 10 most intense precursor ions using stepped normalized collision energies of 30, 50, and 65. Precursor ions were dynamically excluded for 2 s following fragmentation. MS/MS spectra were acquired at a resolution of 30,000 with an AGC target of 1E5.

**Metabolomics and lipidomics data processing**

Raw LC-MS/MS data files from *in vitro* metabolomics and lipidomics experiments were processed using Compound Discoverer v3.3 (Thermo Fisher Scientific). Retention times were aligned across samples using the ChromAlign algorithm[25], followed by peak grouping, gap filling, and peak-area calculation. Peak areas were log10-transformed prior to statistical analysis. Principal component analysis (PCA) was used to visualize patterns in metabolomic profiles. Differences between groups in the PC1–PC2 score space were assessed using permutational multivariate analysis of variance (PERMANOVA) based on Euclidean distances, implemented using the adonis2 function in the R[26] (v4.6.0) package vegan[27] (v2.7-2). Pairwise comparisons were performed between groups, and p-values were adjusted for multiple comparisons using the Benjamini–Hochberg false discovery rate (FDR) procedure Differences in metabolite or lipid abundance among groups were assessed using FDR with Benjamin–Hochberg correction (adjusted p-value < 0.2)

Metabolite annotations were assigned using MS1 and MS2 data based on accurate mass, isotopic pattern, predicted molecular formula, and MS/MS spectral matching against mzCloud, mzVault, ChemSpider, BioCyc, and additional databases integrated within Compound Discoverer. Annotations were considered for downstream analysis when the measured mass was within ±10 ppm of the predicted mass. Metabolites without sufficient MS/MS evidence were considered putative annotations rather than confirmed identifications.

Processed metabolomics and lipidomics data were exported from Compound Discoverer, formatted, and uploaded to MetaboAnalyst v6.0 for statistical and pathway analyses. Data were normalized during data processing as described below. For pathway analysis, metabolites exhibiting an absolute log2 fold change ≥ 1 relative to WT and a raw p-value < 0.05 were selected. Metabolites significantly altered relative to WT, as well as selected lipid species of interest, were visualized using GraphPad Prism version 11.0.2 for Windows. Feature abundances were normalized using the non-background total ion current (TIC) for each sample prior to visualization and comparison.
