## Supplementary material for "A microbiota-dependent fitness locus promotes intraspecies competition and cell envelope resilience in gut bacterium *Phocaeicola vulgatus*": Table Legends

1. **Supplementary Table S1:** Tn-seq transposon library outgrowth bias analysis.
2. **Supplementary Table S2:** Barcodes used for analysis of the mouse fecal and cecal samples that contained the Tn-seq library.
3. **Supplementary Table S3:** Tn-seq data including raw read counts and analysis of genes with Tn insertion.
4. **Supplementary Table S4:** Untargeted metabolomics data showing all identified metabolites and their significance for selected Tn mutants and WT *Pvu*.
5. **Supplementary Table S5:** RNA-seq data including normalized counts, DESeq analysis, and all 14 shared COG category M genes between ∆*difS* and ∆*difH.*
6. **Supplementary Table S6:** Untargeted lipidomics and metabolomics data showing all identified metabolites and their significance for all clean deletion strains compared to WT *Pvu*.
7. **Supplementary Table S7:** ChIP-seq data showing all peaks identified by MACS2 in BHIS and BMM including identifying those with the DNA-binding motif. Additionally, information on the gene neighborhoods of ten up- and downstream genes of each identified peak.
8. **Supplementary Table S8:** All primers used in this work.
